# The EGFR ligand amphiregulin regulates genomic integrity by facilitating heterochromatin formation in response to replication stress

**DOI:** 10.1101/2024.10.27.620431

**Authors:** Tianqi Jiang, Aiman Zein, M.A. Christine Pratt

**Affiliations:** Department of Cellular and Molecular Medicine University of Ottawa, Ottawa, Canada

## Abstract

The EGFR ligand, amphiregulin (AREG) is a key mammary ductal cell differentiation and growth factor. AREG has also been detected in the nucleus of some epithelial cancers although the physiological stimulus and nuclear role are not known. Using immortalized mammary epithelial cells (MECs), we have discovered that AREG undergoes retrograde trafficking to the nuclear membrane (nAREG) in close proximity with lamin A where it is required to both maintain constitutive heterochromatin and transiently increases H3K9me3 in response to replication stress (RS). RS resulted in an increase in AREG protein, enhanced nuclear membrane prelamin A and increased heterochromatin protein, HP1α. In contrast, siRNA-mediated depletion of endogenous AREG reduced HP1α and SUV39h1 proteins accompanied by decompaction and reduction in H3K9me3 heterochromatin despite the presence of soluble AREG. The nuclear membrane (NM) was also impacted resulting in dissipation of the Ran-GTPase gradient, reduced matrix lamin A with increased invaginations. Moreover, AREG knockdown slowed replication fork speed, increased new replication origins and enhanced global transcription while promoting and exacerbating DNA damage in response to RS. DNA damage was most pronounced in AREG-depleted *BRCA2*^mut/+^ MECs which entered senescence following RS, indicating an important nAREG-dependent role in genomic stabilization in these cells. Overall, this study reveals a novel and fundamental role for nAREG in heterochromatin maintenance and the response to RS, that is most critical in *BRCA2*^mut/+^ MECs deficient in replication fork protection.

## Introduction

Amphiregulin (AREG) is a widely expressed EGFR ligand that has pleiotropic roles in differentiation, stem cell maintenance, proliferation, metabolism and inflammation (1). In the mammary epithelium AREG is a key factor controlling normal differentiation, proliferation and ductal elongation (2) as well as stem cell maintenance (3). It is synthesized as a glycosylated plasma membrane-anchored pro-protein and “shedding” stimuli result in cleavage of the ectodomain by the ADAM17 protease. This releases a soluble N-terminal fragment containing an “EGF-like” domain which functions as an autocrine or paracrine EGFR ligand, 4). The level of ectodomain shedding is in part controlled by the endocytosis of mono-ubiquitinated full-length pro-AREG (5) which results in trafficking via the endosomal-lysosome pathway for degradation.

Decades ago, AREG was reported present in the nucleus of ovarian epithelial cells and cancer cells (6) and nuclear localization in fibroblasts was found to be essential for mitogenic activity (7). Unlike EGF, AREG contains an NLS (8) and while AREG shedding releases soluble AREG it also initiates the endocytosis and retrograde trafficking of C-terminal and unshed proAREG to the Golgi then endoplasmic reticulum (ER) followed by nuclear entry via nuclear pores (9). Nuclear proAREG (nAREG) lacks the C-terminal 11aa and interacts with lamin A in the nuclear matrix resulting in increased H2K9me3 heterochromatin (9). The biological stimuli that promotes AREG nuclear trafficking, the underlying mechanism by which AREG regulates heterochromatin and ultimate function are unknown.

Heterochromatin is a repressive histone mark that is either constitutive or facultative. Constitutive heterochromatin is enriched for di– and trimethylated histone H3 lysine 9 (H3K9me2 and H3K9me3 respectively) and represents approximately 50% of the human genome (10). In contrast, facultative heterochromatin forms and disassembles in response to developmental or environmental signals to enable reprogramming of gene expression in a cell lineage specific manner (11). Most constitutive heterochromatin is situated at pericentromeric and telomeric domains of chromosomes where it silences repeats and transposable elements (12) and also dictates the timing of genome compartment replication (13). Perinucleolar regions are enriched for heterochromatin as is the nuclear periphery where it is tethered to the nuclear lamina in regions known as lamina associated domains (LADs) (14,15). Heterochromatin associates with the nuclear lamina directly through heterochromatin protein (HP1α) which binds the lamin B receptor and also via the proline-rich PRR14 protein which interacts with lamin A/C (16,17).

In mammalian cells, heterochromatin is established by families of H3K9-specific histone methyltransferases (HMTs). The suppressor of variegation 3-9 (SUV39)-family proteins are associated with di– and tri-methylation of H3K9 at centromeric and telomeric regions (18). During the replication of constitutive heterochromatin, parental nucleosomes marked with H3K9me2/3 are randomly distributed between the two new copies of DNA. Restoration of me3 modified H3 on newly formed nucleosomes occurs via the recruitment of SUV39h1/2 to the parental histones containing H3K9me2/3 resulting in histone methylation on the new histone (19). SUV39h1 binds to heterochromatin protein 1α (HP1α) (20) which interacts with H3K9me through its chromodomain and can then dimerize and multimerize (21,22), bridging nucleosomes which may help stabilize the nascent heterochromatin domains (23) and resulting in the spread of H3K9me3 over large regions (24).

Lamin A is an intermediate filament integral inner nuclear membrane (INM) protein (25,26). The precursor form, prelamin A, is synthesized in the cytoplasm and interacts on the cytoplasmic side of the ER with SNX6 (27), which is a key protein of the retromer complex that associates with EEA1 early endosomes (28) to mediate retrograde transport of transmembrane cargoes from endosomes to the trans-Golgi network, plasma membrane and ER (29). The nuclear transport of lamin A through Ran-dependent nuclear pore entry is dependent on its association with SNX6 (27). Lamin A undergoes several C-terminal post-translational modifications including farnesylation and methylation followed by cleavage of the prenylated CAAX motif by the ZMPSTE24 metalloprotease. Accumulation of prelamin A forms in *ZMPSTE24*-deficient cells demonstrate increased H3K9me3 heterochromatin as a consequence of enhanced binding of prelamin A with HP1α resulting in its stabilization (30).

Replication stress (RS) resulting in DNA damage is recognized as a significant contributor to genomic instability that can lead to transformation (31). RS can result from numerous impediments to replication fork progression including collisions of the replication-transcription complexes and nucleotide depletion (32). The transient formation of heterochromatin is linked to replication stress (33,34) as well as double strand breaks (35) where formation of heterochromatin serves to recruit repair factors and may reduce transcription to prevent replication:transcription collisions (33). The BRCA2 protein has a key role in preventing DNA damage in response to RS by participating in homologous recombination and replication fork protection (36,37) protecting the reversed fork from excessive degradation by nucleases (38) and ultimately DNA breaks. Thus, *BRCA2*^mut/+^ mammary epithelial cells (MECs) are differentially sensitive to RS-induced DNA damage.

Here, we report our discovery that, in response to RS, AREG undergoes nuclear transport with prelamin A and colocalizes with prelamin A in the INM promoting increased HP1α protein levels accompanied by an increase in H3K9me3 heterochromatin formation. Moreover, AREG contributes to the maintenance of heterochromatin and NM structure as revealed by the depletion of AREG which results in decompaction of heterochromatin, disrupting replication and transcription, precipitating DNA damage and the formation of NM invaginations. Overall, through maintenance of constitutive heterochromatin and mediating a transient increase in heterochromatin in response to RS, we conclude that nAREG helps maintain genomic stability, a function critically important in *BRCA2*^mut/+^ MECs where chronic DNA damage as a consequence of RS could lead to senescence.

## Methods

### Cell culture

hTERT mammary epithelial cells were obtained from Dr. C. Lewis (UT Southwestern, TX and were established as described (39). Cell are maintained in DMEM/F12 (Wisent, 319-075-CL) containing 0.4% pituitary extract (Gibco, 13028-014), Glutamax (Gibco, 35050-061), amphotericin B, gentamicin, penicillin/streptomycin, supplemented with 5ng/ml EGF, 0.5ug/m hydrocortisone, 5ug/ml insulin, 5mM 3,3,3-tri-iodo-thyronine and 5nM isoproterenol (all Sigma) incubated at 37°C humidified atmosphere under 5% CO_2_. For all siRNA transfections, 15nM of soluble AREG was added to the medium to obviate EGFR-mediated effects in AREG-depleted MECs. For nucleoside supplementation, EmbryoMax® Nucleosides 100× (ES-008-D, Sigma-Aldrich) were used with 50-fold dilution.

### siRNA reverse transfection

hTERT-MECs were grown for 72h before transfection to ensure optimal growth period. On the day of transfection, a mixture of siRNA (Dharmacon ON-TARGETplus SMART Pool human AREG siRNA or Non-targeting pool, Horizon (D-001810-10-05) and transfection reagent (Dharmacon T-2001-01) was made by diluting with hTERT media. The mixture was then incubated in 37°C for 20 mins. The hTERT-MECs were then trypsinized, collected, and combined with the transfection mixture. The cell number was adjusted according to the company’s protocol. hTERT-MECs were transfected for ∼60h before switching into fresh media for 12 hours prior to harvest or treatments.

### Construction of lenti-AREGΔC11

Lenti-AREG (C-terminal V5 tag) was purchased from GeneCopia (EX-OL00093-LX304). The C-terminal 11 amino acids were deleted using the Q5® Site-Directed Mutagenesis Kit (New England Biolabs, E0554S). Primers were designed using the NEBase Changer tool to generate the deletion were: forward-3’-TACCCAACTTTCTTGTAC-5’ and reverse-3’-TACCCAACTTTCTTGTAC-5’. Plasmids were verified by sequencing.

### DNA comet assays

Assays were performed as described in neutral conditions and alkaline conditions (100,101). Microscope slides were coated with 1% LMP agarose (FroggoBio, A87-500G). Prepared slides were let cooled at room temperature overnight. The next day, cells were trypsinized and collected in PBS on ice. 1% LMP agarose (Bio-Rad, 161-3111) kept at 37°C was mixed with collected cells at ∼1000 cells/ul. Around 75ul of cell mixture was dispensed onto the coated slides and flattened with a coverslip. The slides were then chilled on ice to allow the agarose to harden. Subsequently, another layer of LMP agarose was coated onto the sample. Coverslip is then removed and slides containing the cells were placed into lysis buffer (2.5M NaCl, 100mM EDTA, 10mM Tris-base, pH 10, in ddH2O) at 4°C for at least 1h. The following day, slides were carefully washed with Tris buffer (0.4M Tris, pH 7.4, in ddH2O) before placing into electrophoresis chamber. For alkaline comet assay, electrophoresis buffer was made using 10N NaOH and 200mM EDTA at pH 13. For Neutral comet assay, electrophoresis buffer was made using 300mM sodium acetate and 100 mM Tris-HCl at pH 8.3. Voltage was set to ∼0.74 V/cm of chamber length and electrophoresis was performed. SYBR GOLD staining (labels ssDNA and dsDNA) was performed and slides containing the gel was let dry before immediately imaging. A neutralization (0.4M Tris, pH 7.4, in ddH2O) washing step was performed prior to akaline comet assay staining.

### DNA fiber assay

Assays were performed essentially as described (42). Thymidine analogs solution: 2.5 mM 5-chloro-2′-deoxyuridine (CldU) stock solution and 2.5 mM 5-iodo-2′-deoxyuridine (IdU stock solution were prepared in DMEM/F12. CldU was vortex at room temperature until dissolved and IdU was vortexed after incubated at 60°C. All analogs were aliquoted and stored at –20°C until used. Cells were plated in 6-well plate at 300,000 cells per well and treated with siAREG and siNT for 72h. CldU and IdU were mixed with hTERT media at 25uM and 250uM respectively and incubated for 20min each. HU (5mM) was added subsequently to CldU or IdU to investigate its effect on fork restart and protection, respectively. After incubation, cells were trypsinized with TrypLE and centrifuged for collection. Each sample was then resuspended in ice-cold 1XPBS at 500,000 cells/ml and 2ul of cell suspension was spotted on the silane-coated slide (Sigma-Aldrich, S4651). After 5mins of room temperature incubation, lysis buffer (31.52mg/ml Tris hydrochloride, 14.6mg/ml disodium EDTA, 5mg/ml SDS in ddH2O) was added onto the cell suspension and incubated for another 10mins at room temperature. The slides were then tilted at 25° to 40° to allow the suspension to run down (estimated 5mins); the suspension was air-dried completely. The slides were fixed with 3:1 methanol/acetic acid fixature at –20°C for 15mins and incubated in 70% ethanol at 4°C overnight. The next day, the slides were incubated in 2.5M HCl at 37°C for 1h, washed with 1XPBS, and then blocked with 0.5% BSA in 1XPBS for 30mins. The slides were then incubated with 1:300 monoclonal anti-BrdU antibody [BU1/75 (ICR1)] (Abcam, ab6326) and 1:300 Mouse monoclonal anti-BrdU antibody (clone B44) (Becton Dickson, 347580 (7580)). The slides were then fixed with 4%PFA in room temperature for 15mins and washed with 1XPBS. Next, the slides were incubated with 1:400 Alexa Fluor 555 goat anti−rat IgG (Thermo Fisher Scientific, A21434) and 1:400 Alexa Fluor 488 F (ab′)2 goat anti−mouse IgG (Thermo Fisher Scientific, A-11017). These slides were washed with 1X PBS and mounted with Vectashield plus antifade mounting medium (Biolynx, VECTH19002) for imaging.

### Proximity Ligation Assay (PLA)

hTERT-MECs were fixed with 4% paraformaldehyde (PFA) for 15 minutes at room temperature and then washed 3 x 5mins with PBS. hTERT-MECs were permeabilized 0.3% Triton-X 100 in PBS for 30 minutes. After washing with buffer A (Millipore, DUO82049), the cells were blocked with provided blocking solution (Millipore, DUO82007) for 1 hour at room temperature. Primary antibodies of interest were diluted in Duolink antibody diluent (Millipore, DUO82008) and incubated with the cells at room temperature for 1h or alternatively overnight at 4°C. Subsequently, cells were washed 3×5mins with buffer A before incubation with PLA anti-rabbit PLUS (Millipore, DUO92002) and anti-mouse MINUS probes (Millipore, DUO92004) at 37°C for 1 hour. After washing off the unbound probes, ligation was performed using the ligation kit (Millipore, DUO92008) at 37°C for 30 minutes, allowing the formation of circular DNA templates if the primary antibodies targeting the proteins of interests were in close proximity. This was followed by the addition of polymerase which amplifies and adds the fluorescently labelled, complementary oligonucleotide probes. Finally, cells were counterstained with DAPI for 10 minutes and mounted with antifade mounting medium for imaging.

### Image analysis

Images were analyzed using Fiji (Image J) open source software or Imaris Image analysis software. To quantify lamin A thickness, we utilized the “Local Thickness plugin” in Fiji (https://imagej.net/Local_Thickness). For further analysis, the thickness measurement distribution was quantified using the histogram option (Analyze > histogram). Comet tail moments were quantified using the OpenComet plugin in Fiji (https://cometbio.org/)

### Statistical analysis

Results were expressed as mean ± S.E.M. of a minimum of 100 cells per treatment group. Two-tailed Student’s t-test and one-way ANOVA analyses were performed using GraphPad Prism software. A minimum p-value < 0.05 was considered statistically significant.

**Additional methods** can be found in Supplementary files.

## Results

### Replication stress increases AREG peptide levels and nuclear membrane localization with lamin A

Isokane and colleagues (9) demonstrated that treatment of HeLa cells with TPA results in retrograde nuclear trafficking of AREG and nuclear localization. Both phorbol esters such as TPA (45) and DNA damage can activate the stress kinase, p38 (46). We therefore used hTERT-immortalized MEC isolates derived from two normal and two BRCA2^mut/+^ hTERT-immortalized primary MECs (43), to analyze the effects of RS on AREG protein and localization. MECs were treated with hydroxyurea (HU) (4mM for 4hrs) followed by immunofluorescence (IF) for AREG and lamin A. Untreated MECs show the typical distribution of AREG, primarily at the plasma membrane (Figure 1A). In contrast, HU-treated MECs contained increased nAREG which was randomly colocalized at and within the inner NM with lamin A. Multiple fragments of soluble and intracellular AREG have previously been identified in different types of epithelial cells (9, 44, 45). Similar to the array of AREG peptides found in breast cancer cell lines (49), immunoblots showed that both normal and *BRCA2*^mut/+^ MECs expressed baseline levels of 37kDa-40kDa AREG and AREG and additional bands of 23kDa, 18kDa and 16kDa. The AREG antibody recognizes regions from aa107-184 indicating that all peptide bands contained sequences within the EGF-like domain and C-terminus (1,4). Notably, RS resulted in a marked increase in the 23kDa and 18kDa species and a diminution of the 16kDa band suggesting that the latter is likely derived from these larger peptides. This was most profound in *BRCA2*^mut/+^ MECs and increased up to 3h following washout of HU (Figure 1B). HU resulted in an approximate 2-fold increase in AREG mRNA (Figure 1C) which could contribute to an increase in total AREG protein.

**Figure 1.**
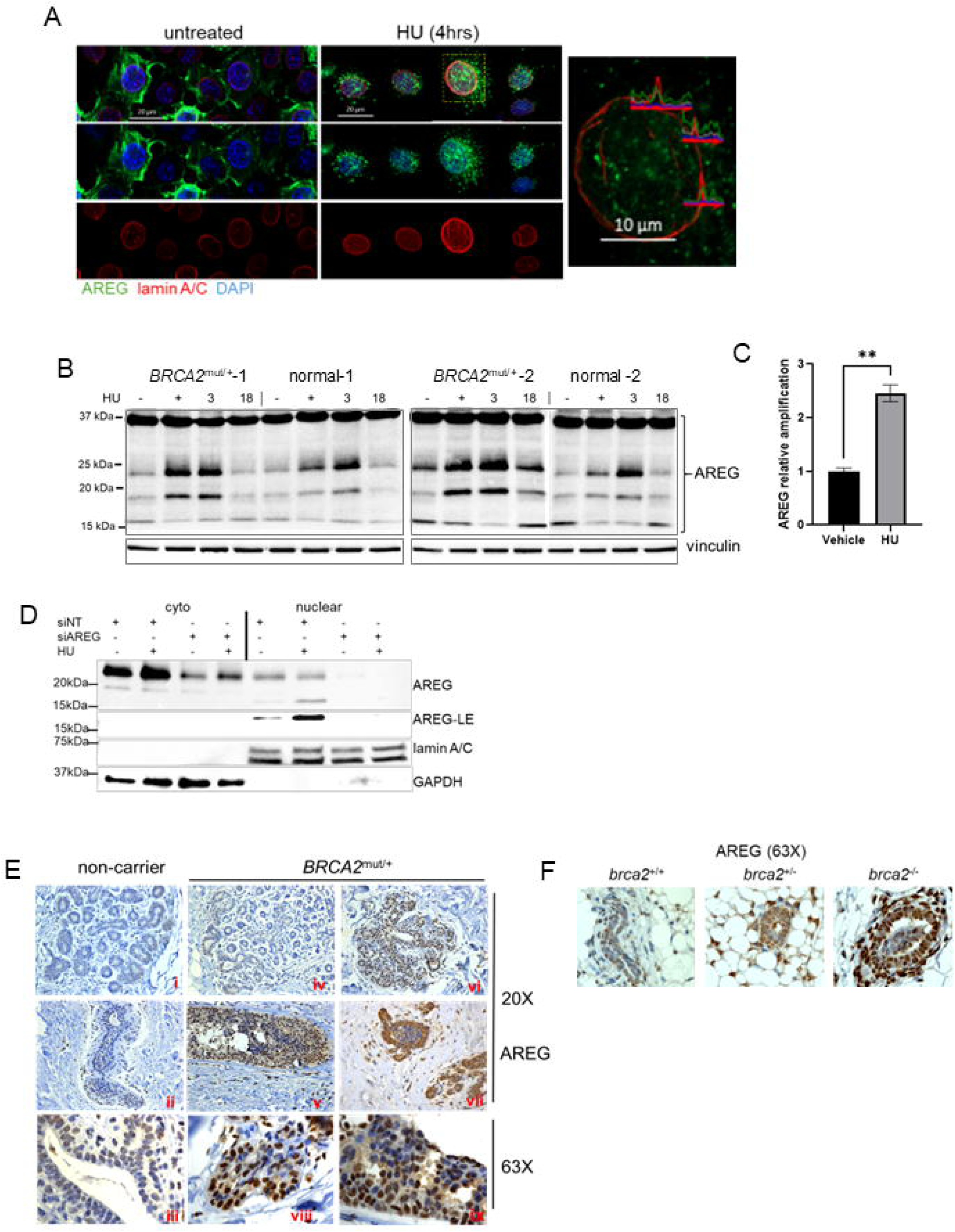
Nuclear AREG expression correlates with mouse mammary Brca2 deficiency and dysplastic regions of *BRCA2*^mut/+^ human mammary tissue and is induced by replication stress in MECs. **A**, Immunofluorescence of AREG and lamin A in *BRCA2*^mut/+^ MECs treated with vehicle and 4mM HU for 4h. Bars are 20µm. The boxed area is magnified on the right panel showing overlap in IF intensity for AREG and lamin A at regions along the NM. Bar is 10µm. **B,** Immunoblot for AREG in *BRCA2*^mut/+^ and normal human hTERT-MECs treated with vehicle (H_2_O) or 4mM HU for 4 h (HU), followed by replacement of fresh media without HU and collection at 3h and 18h. **C,** Graph of qPCR to detect AREG mRNA in untreated, HU-treated and MECs 3h after change to media without HU. Bars are S.E.M. of 3 independent experiments in triplicate. **D,** Immunoblot of nuclear and cytoplasmic fractions from MECs transfected with siAREG or the siNT control. 72hrs after transfection MECs were treated with vehicle or 4mM HU for 4h and harvested then lysates probed with antibodies to AREG, lamin A and GAPDH. LE –long exposure of the nuclear 18kDa band. Data shown is representative of at least 3 independent experiments. **E,** Examples of AREG IHC in patient mammary glands from non-carriers (i-iii) and *BRCA2*^mut/+^ mammary glands (iv-ix). A range of nAREG expression was present in histologically normal tissue from both patient groups (i-iv). nAREG was detected in the majority of *BRCA2*^mut/+^ MECs in regions of mild dysplasia and hyperplastic ducts (v-vii). Higher magnification in viii and ix show nAREG-positive cells in the luminal layer. Images are representative of findings from 10 normal and 10 *BRCA2*^mut/+^ glands. **F,** AREG IHC in mammary gland tissue from MMTV-cre transgenic mice with *brca2^+/+^*, *brca2^+/f^* or *brca2^f/f^*alleles shows detection of nAREG in *brca2^-/-^* mammary ducts. Moderate to negative AREG staining was observed in *brca2^+/+^* and ^-/-^ glands.

Using antibody against full length AREG, we next assessed the effect of acute RS on the subcellular location of AREG in cytoplasmic and nuclear fractions in *BRCA2*^mut/+^ MECs with or without RS (Figure 1D). In control conditions, the 23kDa AREG was present in the cytoplasm and to lesser extent was also found in the nuclear fraction. While the 23kDa peptide did not change following HU, the 18kDa nuclear species was increased, consistent with an overall increase in nAREG. Thus, intracellular AREG undergoes cleavage and a constitutive level is present in at least some nuclei of unperturbed MECs while RS increases nuclear residence of AREG peptides.

The presence of nuclear-localized AREG in cancer and normal cells has been documented in several studies (6,47,48). To investigate this further in the context of normal and high risk mammary epithelium, we performed AREG IHC on mammary tissue sections from normal reduction mammoplasty and *BRCA2*^mut/+^ prophylactic mastectomy, which we hypothesized would have greater levels of nAREG based on increased endogenous RS. Regions of *BRCA2*^mut/+^ histologically normal tissue largely expressed similar levels of AREG as tissue from *BRCA2* normal patients (compare panels i, ii and iv), although some areas, including both lobules and ducts, were strongly positive for nAREG in *BRCA2*^mut/+^ sections (Figure 1E; panels vi and viii). Breast tissue from *BRCA1/2* mutation carriers is characterized by increased incidence of hyperplastic and premalignant lesions (49,50). Notably regions of dysplastic and overt hyperplasia in *BRCA2*^mut/+^ tissue with intense nuclear localization of AREG in up to 90% of cells were detected (panels v, vii and ix). As expected, some stromal cells, most likely fibroblasts, also stained with anti-AREG (blue arrowheads). Thus nAREG is most associated with MECs likely to undergo RS and DNA damage. We further determined if nAreg correlated with *Brca2* status in mouse mammary tissue. IHC on glands from 6-week old MMTV-cre;*Brca2*^+/+^; *Brca2*^f/+^ and *Brca2*^f/f^ mice showed that mammary epithelium from both wt and *Brca2*^f/+^ mice expressed low to moderate levels of Areg, however prominent nAreg staining was present in a the majority of *Brca2*^f/f^ MECs, potentially a result of chronic RS due to the absence of Brca2 (Figure 1F).

### Replication stress induces p38-dependent AREG trafficking with lamin A and nuclear periphery localization

AREG was reported to interact directly with lamin A as determined by immunoprecipitation (9). To further determine the impact of RS on the nuclear trafficking of AREG with respect to lamin A, we performed antibody-primer rolling circle amplification proximity ligation analysis (PLA) which detects two proteins localized within a distance of <40nm. Using AREG and lamin A antibodies, a low number of PLA foci were present in untreated cells, however foci were significantly increased following the 4 hr treatment with HU. AREG:lamin A foci were found in the cytoplasm (ER) as well as at the nuclear periphery and nucleoplasm (Figures 2A, 2B). The increase in nAREG in response to 4hr of HU-induced RS requires rapid nuclear trafficking over this time period. These results suggest that, in response to RS, lamin A on the cytoplasmic surface of the ER (27) may interact with endosome-delivered AREG giving rise to the significant increase in lamin A:AREG PLA foci observed in HU-treated MECs.

**Figure 2.**
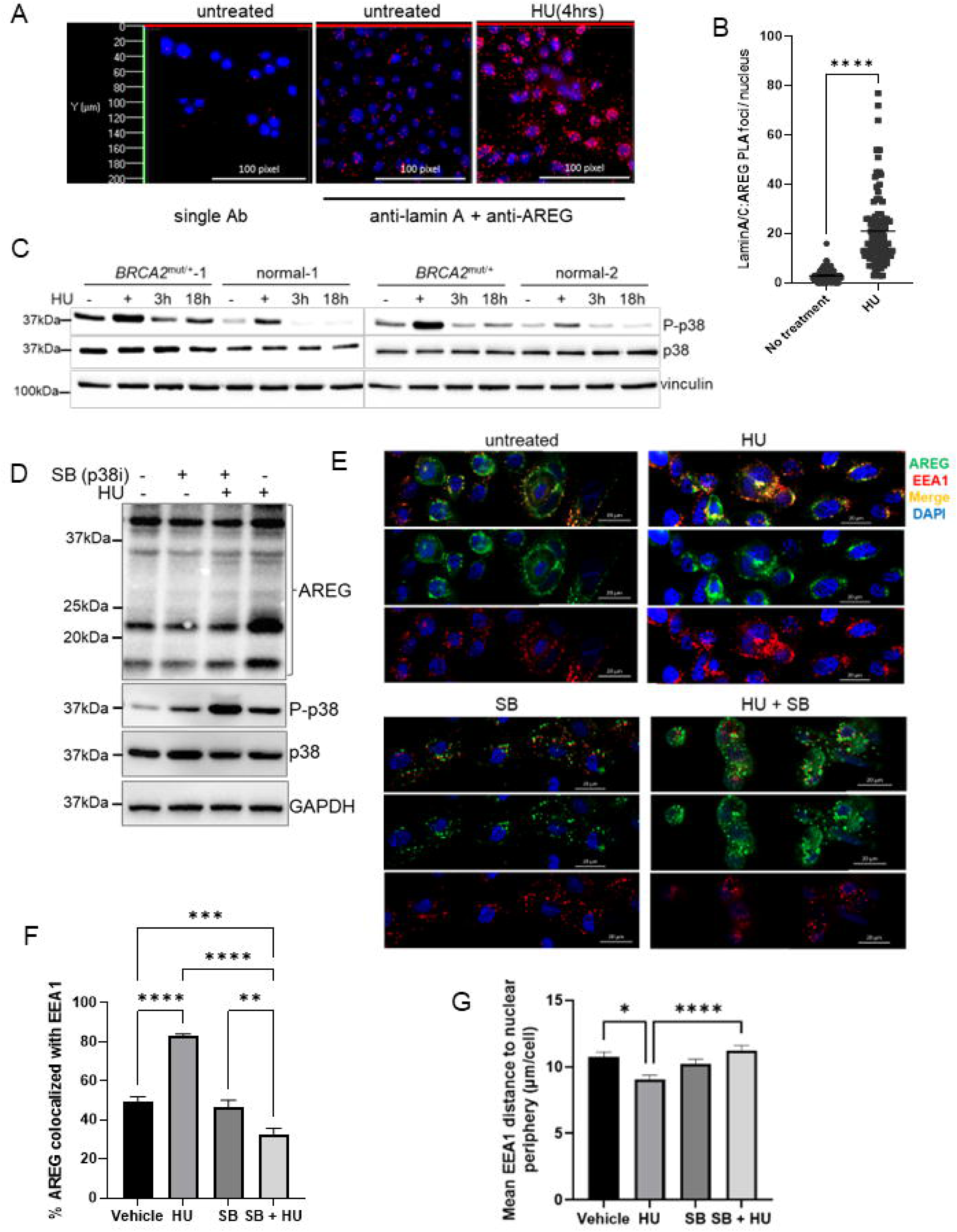
Replication stress increases cytoplasmic and nAREG:lamin A interaction and AREG association with early endosomes. **A**, PLA fluorescence analysis of AREG: lamin A interaction in *BRCA2*^mut/+^ MECs treated with vehicle or HU for 4h. Images were acquired on a Zeiss AxioObserver-7 microscope and rendered in 3D images using Zeiss Zen Blue software (35 z-stacks). A single antibody control is also shown. **B,** Analysis of foci numbers per nucleus corrected for single antibody foci were calculated using Imaris software. Data represent analysis of 100 nuclei per condition from 3 independent experiments. **** p<.0001; *p<.01 (one-way ANOVA). **C,** Immunoblot of P-p38α in lysates from *BRCA2*^mut/+^ MECs and normal MECs following HU-induced RS. **D,** Immunoblot of lysates of *BRCA2*^mut/+^ MECs treated for 4h with vehicle, 10µM SB208530, 4mM HU or 4mM HU with 10µM SB and probed with antibodies for AREG, P-p38α and total p38α. **E,** Dual IF for AREG and EEA1 on *BRCA2*^mut/+^ and normal MECs treated as in D. Bars are 20µm. **F,** Graph of Imaris analysis of colocalization of EEA1 and AREG. Data represent analysis of ≥100 cells per group from 3 independent experiments. **G,** Graph of the average distance between the nuclear periphery and EEA1 foci in MECs treated as indicated. Each average is from a minimum of 50 cells per group. For both graphs *p<.05 **p<.01; ***p<.001; ****p<.0001 One-way ANOVA.

As discussed, DNA damage signaling activates p38 (46) which can stimulate endocytosis (54) and phosphorylate EEA1 to regulate recruitment to membranes (55). Consistent with greater DNA damage, P-p38α was strongly induced by HU in *BRCA2*^mut/+^ MECs compared to normal MECs (Figure 2C) and activation was commensurate with the increased level of AREG protein shown in Figure 1. We therefore tested the effect of p38 inhibitor SB203580 (SB) on the induction of AREG following HU treatment. Note that SB does not decrease P-p38α but prevents substrate phosphorylation (56). Figure 2D shows that SB prevented the HU-mediated increase in AREG protein. IF of EEA1 positive early endosomes and AREG in Figure 2E revealed many cells with predominantly plasma membrane AREG and scattered EEA1 foci. AREG and EEA1 were also colocalized at the nuclear periphery in some cells. HU strongly increased nuclear periphery colocalization of AREG-associated with EEA1endosomes. AREG became localized in clumps at the plasma membrane and throughout the cytoplasm with SB treatment alone. Although AREG was more diffuse than SB only treated MECs, the co-addition of HU and SB prevented the nuclear periphery accumulation of AREG. Together these results are consistent with RS-induced DNA damage activation of p38α in MECs which promotes early endosome trafficking of AREG and AREG:lamin A nuclear entry. Analysis in Fiji showed the increase in colocalization of AREG and EEA1 after HU treatment and inhibition by SB (Figure 2F). Figure 2G depicts the distances between EEA1 foci and DAPI at the nuclear periphery under all conditions and indicates that HU significantly increases nuclear proximity of EEA1 foci which is prevented by SB. These results are consistent with redirection of endosomal AREG toward the nucleus during RS.

### Endogenous nAREG promotes H3K9me3 formation, increases HP1α protein levels and alters prelamin A localization

Previous studies showed that enforced expression of AREG lacking the C-terminal 11 amino acids results in NM localization where it interacts with lamin A and enhances H3K9me3 heterochromatin (9). Since AREG undergoes significant nuclear trafficking following HU treatment, we next analyzed the formation of H3K9me3 after HU in siAREG vs siNT-transfected MECs. Immunoblot analysis in Figure 3A revealed that, as expected, HU transiently increased H3K9me3, however, AREG KD strongly reduced both baseline and HU-induced transient H3K9me3.

**Figure 3.**
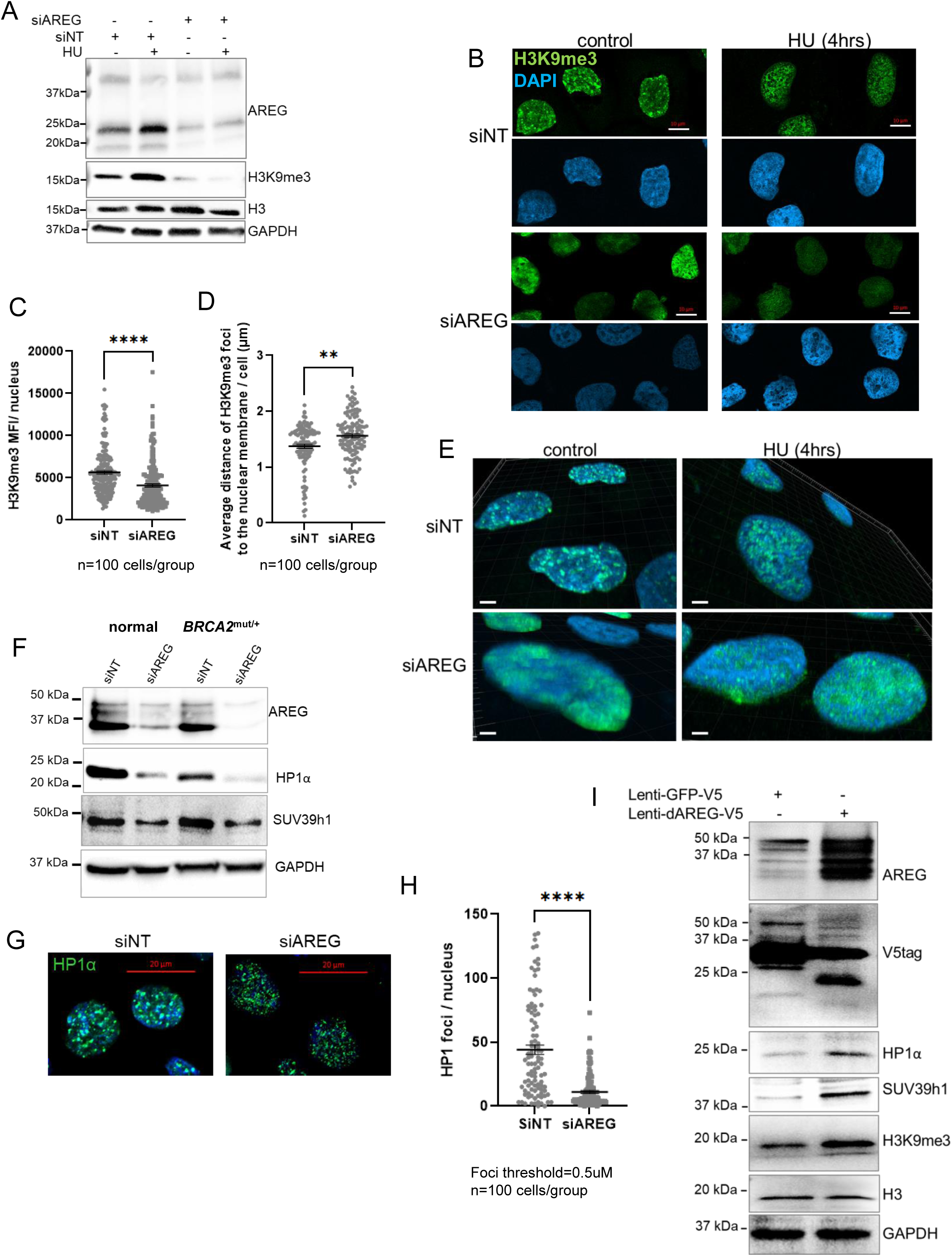
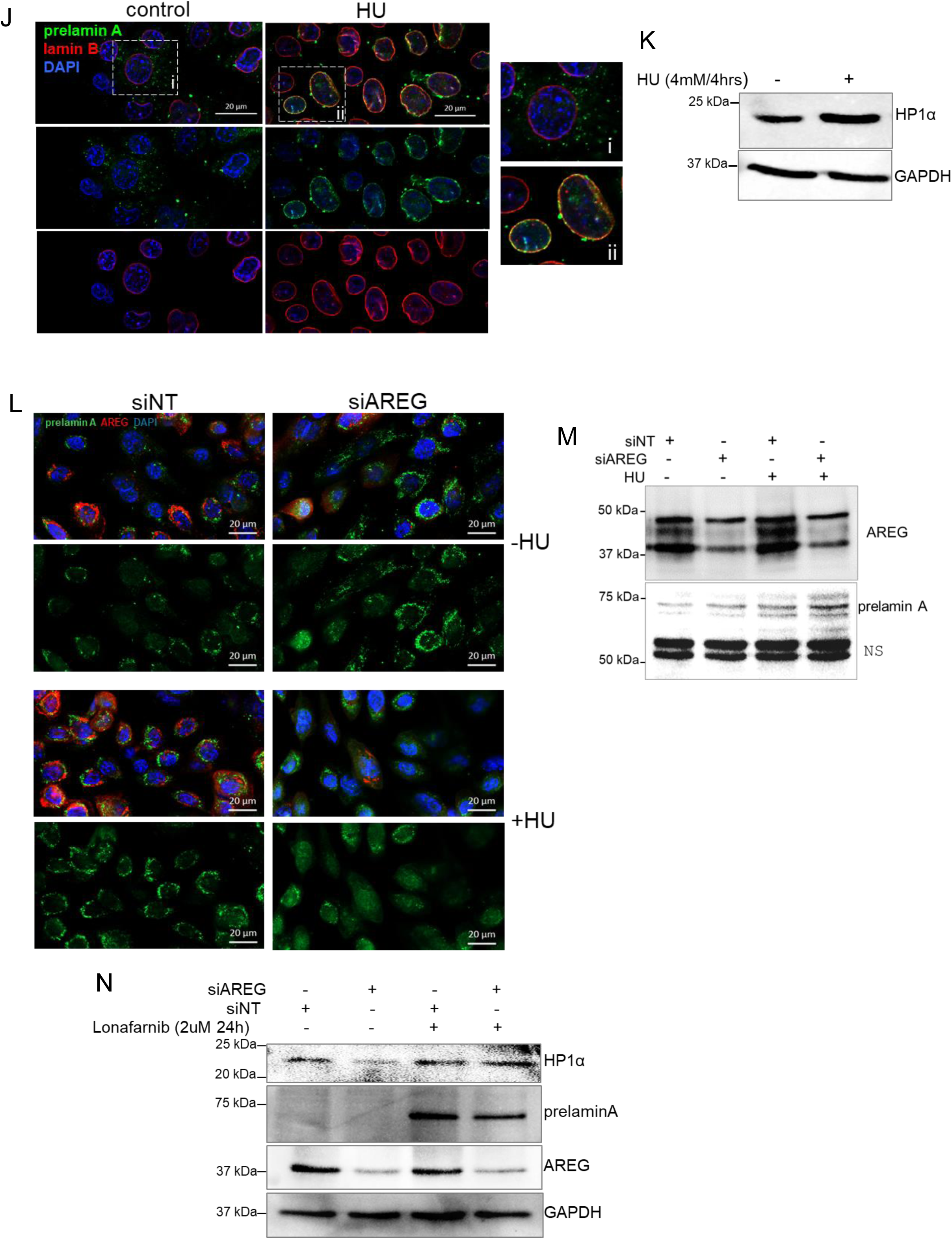
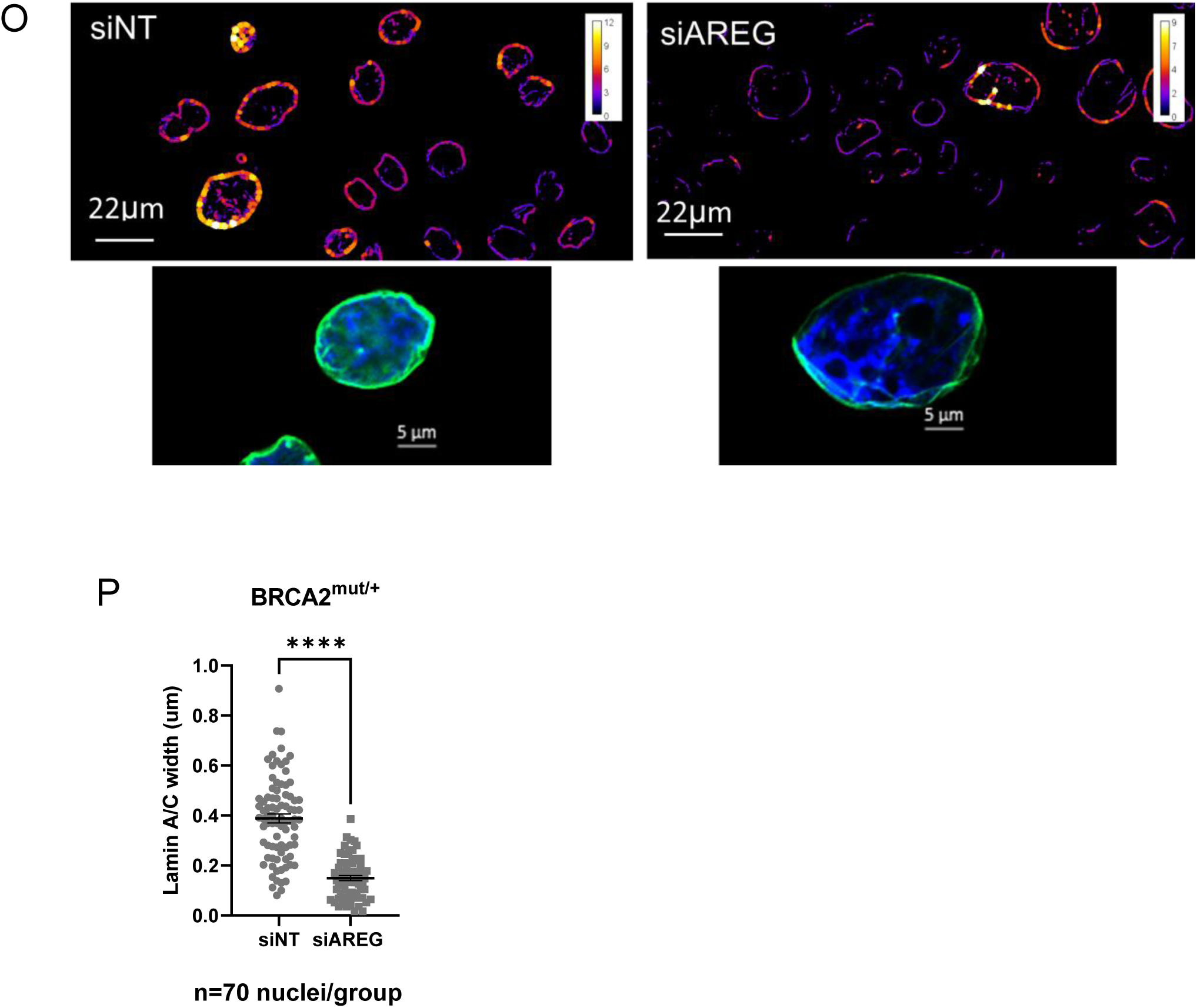
Replication stress and nuclear trafficking of AREG promote H3K9me3 increase nuclear prelamin A residence. **A**, Representative immunoblot analysis (n=4) of AREG and H3K9me3 and histone H3 in *BRCA2*^mut/+^ MECs treated with vehicle or HU for 4h. **B,** Confocal images of H3K9me3 IF and DAPI in *BRCA2*^mut/+^ MECs 72h following transfection with siNT control or siAREG and treated with vehicle or HU for 4hrs. Bars are 10 µm. **C,** Graph of H3K9me3 mean fluorescence intensity showing the reduction following AREG depletion. n=200 nuclei/ ****p<.0001; two-tailed t-test. **D,** Graph showing reduction of perinuclear H3K9me3 as assessed by distance of H3K9me3 IF from DAPI-defined perimeter performed using Fiji (n=100 cells/** p<.01). **E,** 3D Imaris rendering of H3K9me3/DAPI in cells treated as indicated. **F,** Representative blot (n=2) of normal and *BRCA2*^mut/+^ MECs transfected with siNT or siAREG and 72hr later lysates collected and probed with antibodies for AREG, HP1α, SUV39h1 and GAPDH to control for protein loading. **G,** IF of HP1α depicting chromocentres in siNT– and siAREG-transfected MECs. **H,** HPα marked chromocenters defined as foci of >0.5µm were quantified using Fiji software. Bars are 20um. (n=100 cells per group/****p<.0001; two-tailed t-test). **I,** *BRCA2*^mut/+^ MECs infected with lenti-GFP-V5 or lenti-AREGΔ11-V5 were probed with antibodies to AREG, V5, HP1α, SUV39h1, H3K9me3 and H3. Anti-GAPDH was used to control for protein loading. **J,** IF of prelamin A and lamin B in untreated control MECs and after a 4hr 4mM HU exposure. Magnification of boxed areas are shown on the right. **K,** Representative immunoblot for HP1α in *BRCA2*^mut/+^ MECs treated with vehicle or HU as indicated. GAPDH was used as a protein loading control. (n=2 independent experiments) **L,** IF of prelamin A and AREG in siNT and siAREG transfected MECs treated with vehicle (-HU) or 4mM HU for 4 hrs. **M,** Immunoblot for AREG and prelamin A in MECs treated as indicated. **N,** Immunoblot detection of HP1α in *BRCA2*^mut/+^ MECs transfected with siNT or siAREG and treated vehicle or lonafarnib. **O,** Analysis of lamin A thickness in response to AREG KD. The heatmap on the right indicates thickness. Examples of lamin A IF (green) in MECs from each condition are shown below. **P,** Lamin A thickness was analyzed in Fiji. n=70 cells per group. ****p<.0001 t-test.

In unchallenged cells, Ataxia Telangiectasia-rad related (ATR) kinase-deficiency enhances RS and aberrantly increases perinuclear heterochromatin (54,55). To determine if the nuclear trafficking of AREG and formation of H3K9me3 is induced by an alternate RS stimulus, using the ATR inhibitor, VE-821, we tested the effect of inhibition of on basal and HU-induced AREG protein and H3K9 heterochromatin formation (Supplementary Figure S1A). As expected, HU robustly induced H3K9me3, however the combination of VE-821 and HU blocked this response. Additionally, the increase in AREG protein was also prevented. ATR inhibition also reduced nAREG detection by IF in MECs treated with HU (Supplementary Figure S1B). Thus, ATR activation contributes to the RS-induced increase in AREG protein as well as H3K9me3 formation. In contrast, VE-821 treatment in unchallenged *BRCA2*^mut/+^ MECs, similar to HU, strongly enhanced AREG protein detection in the ER and at the NM while increasing both AREG protein and H3K9me3 levels. Overall, these data demonstrate that endogenous AREG regulates H3K9me3 heterochromatin and extend these findings to reveal that RS originating via different mechanisms can function as a stimulus for AREG nuclear trafficking.

HP1α and lamin A are involved in tethering heterochromatic lamina associated domains (LADs) at the nuclear periphery (16,17). To determine the effect of AREG depletion on global heterochromatin organization, we performed confocal imaging of H3K9me3 IF in *BRCA2*^mut/+^ MECs transfected with AREG or NT siRNAs and treated with or without HU (Figure 3B). In siNT-transfected MECs, H3K9me3 was located at the nuclear periphery and in punctate domains within the nucleoplasm. In contrast, in a majority of AREG-KD cells, overall H3K9me3 mean fluorescence intensity (MFI) was strongly reduced (Figure 3C) or displayed highly diffused H3K9me3 with reduced H3K9me3 localization at the nuclear periphery (Figures 3B, 3D). HU treatment in control MECs resulted in redistribution of H3K9me3 to the nuclear interior in a honeycomb pattern, while H3K9me3 in AREG-depleted MECs remained low and diffuse following HU treatment. 3D images of H3K9me3 in MECs were generated using Imaris software (Figure 3E). Perinuclear and nucleoplasmic foci were evident in siNT-transfected MECs, while HU induced the formation of nucleoplasmic foci. In contrast, AREG-depleted MECs heterochromatin decompaction and/or reduced H3K9me3 while large numbers of small puncta formed in response to HU.

Localized H3K9me3 binding by HP1α is proposed to promote expansion of H3K9me3 on surrounding nucleosomes by recruiting SUV39h1 (20) while HP1α binding to SUV39h1 also results in its stabilization (56,57). Therefore we assayed the effect of AREG depletion on these proteins. Immunoblot analysis of HP1α and SUV39h1 proteins in siAREG-transfected MECs showed that reduction of AREG decreased both HP1α and SUV39h1 (Figure 3F). Heterochromatin bound HP1α has been observed to form collapsed globules which achieve segregation of domains called chromocentres (61). Consistent with reduction of HP1α, large HP1α-marked foci were significantly reduced by AREG depletion (Figures 3G, 3H). To further assess the effect of nAREG on these proteins we infected cells with a lenti-virus expressing an AREG C-terminal 11 amino acid deletion (AREGΔC11) previously shown to traffic to the NM and promote a global increase in heterochromatin (9) or a control GFP-V5 lentivirus. The immunoblot in Figure 3I shows that AREGΔC11 increased cellular levels of HP1α, SUV39h1 and H3K9me3.

Given the association of AREG with cytoplasmic lamin A we reasoned that AREG might regulate processing and/or nuclear transport of prelamin A. Newly synthesized prelamin A is associated with the outer ER prior to entry into the nucleus via Ran-GTPase-dependent entry via nuclear pores. Farnesyltransferase and ZMPSTE24 are present both at the ER and in the nucleus and evidence suggests that most prelamin A may be farnesylated in the nucleus facilitating incorporation into the nuclear matrix where it is rapidly processed via ZMPSTE24-mediated cleavage to yield mature lamin A (59,60). We first considered the effects of RS on nuclear prelamin A in HU-treated, untransfected MECs. Dual IF was performed using an antibody specific for the amino acids C-terminal to the cleavage site for ZMPSTE24 (clone PL-1C7) and anti-lamin B. Figure 3J shows that, in most cells, prelamin A was not detected or weakly detected in the nuclear lamina indicated by lamin B in untreated cells. In contrast, HU resulted a majority of cells exhibiting prelamin A colocalized to the NM with lamin B indicating robust incorporation of prelamin A into the inner NM in response to RS/DNA damage. Importantly, HU-induced RS alone resulted in a small but reproducible increase in HP1α (Figure 3K and Figure S2) similar to that observed in *ZMPSTE*^-/-^ cells (30).

To assess the role of nAREG, prelamin A and AREG IF was performed in MECs transfected with siNT or siAREG and treated with or without HU. The deconvoluted z-stack images in Figure 3L show that, similar to untransfected MECs, prelamin A is detected at the NM of some siNT-transfected cells while in other cells it was colocalized with AREG at least partially in the cytoplasm/ER. A subset of cells were devoid of detectable prelamin A. As expected, HU stimulated an increase in AREG in the perinuclear region and nuclear rim associated with prelamin A. Although some prelamin A was present at the NM after AREG KD, prelamin A was detected predominantly in a cytoplasmic location, likely at the ER. HU-induced RS resulted in diffuse nucleoplasmic staining. Interestingly, immunoblot of cells treated as indicated shows that HU increased prelamin A most strongly in AREG-depleted MECs (Figure 3M). Given the increase in nucleoplasmic prelamin A this increase may be due to nuclear accumulation of non-farnesylated prelamin A since this distribution is reminiscent of cells treated with farnesyltransferase inhibitors (64). To determine the role of prelamin A in AREG regulation of HP1α protein levels, we treated MECs with the farnesyltransferase inhibitor, lonafarnib which strongly increases levels of prelamin A. Immunoblot analysis revealed that lonafarnib prevented the reduction in HP1α in cells depleted of AREG (Figure 3N).

Prelamin A accumulation outside the nucleus in AREG KD MECs could be anticipated to reduce INM lamin A. Indeed, measurement of IF intensity in the INM demonstrated a significant decrease in membrane lamin A in siAREG-transfected MECs (Figures 3O, 3P). To further verify that nuclear-localized AREG promotes prelamin A nuclear accumulation, we infected MECs with either lenti-AREGΔC11-V5 or lenti-GFP-V5 and 24hrs later performed IF. In control MECs expressing lenti-GFP, low levels of AREG and scarce, heterogeneously distributed prelamin A were detected. High levels of AREGΔC11 expression corresponded to strong perinuclear AREG staining and prelamin A was readily detected at the nuclear periphery and nucleoplasm (Supplementary Figure S3).

Together these observations are consistent with AREG:prelamin A nuclear trafficking in MECs which increases HP1α protein levels to maintain constitutive heterochromatin and promote a transient increase in H3K9me3 formation in response to RS.

### Loss of endogenous AREG results in genomic instability

Both the presence of heterochromatin and loss of heterochromatin present challenges to DNA repair and replication. Therefore, we next investigated the effects of AREG KD on DNA damage in MECs. We first evaluated the effect of AREG depletion on combined DNA breaks (SSBs and DSBs) using alkaline comet assays in both normal and *BRCA2*^mut/+^ MECs. Figures 4A shows representative comets from normal and *BRCA2*^mut/+^ MECs treated as indicated and quantified in Figure 4B. While no significant increase in DNA damage was obtained after AREG depletion alone relative to control, HU significantly increased the Olive tail moment in siNT-transfected normal MECs which was further amplified in siAREG transfected MECs. *BRCA2*-insufficiency can result in ssDNA gaps (65) which are exacerbated after treatment with HU (62,63). In keeping with this, HU strongly increased DNA breaks in control transfected *BRCA2*^mut/+^ MECs. In contrast to normal cells, significantly more frequent and extensive DNA damage was evident after AREG depletion in these MECs which was further amplified following HU treatment. These results indicate that endogenous AREG protects MECs from RS-induced DNA damage in MECs which is accentuated in *BRCA2*^mut/+^ MECs.

**Figure 4.**
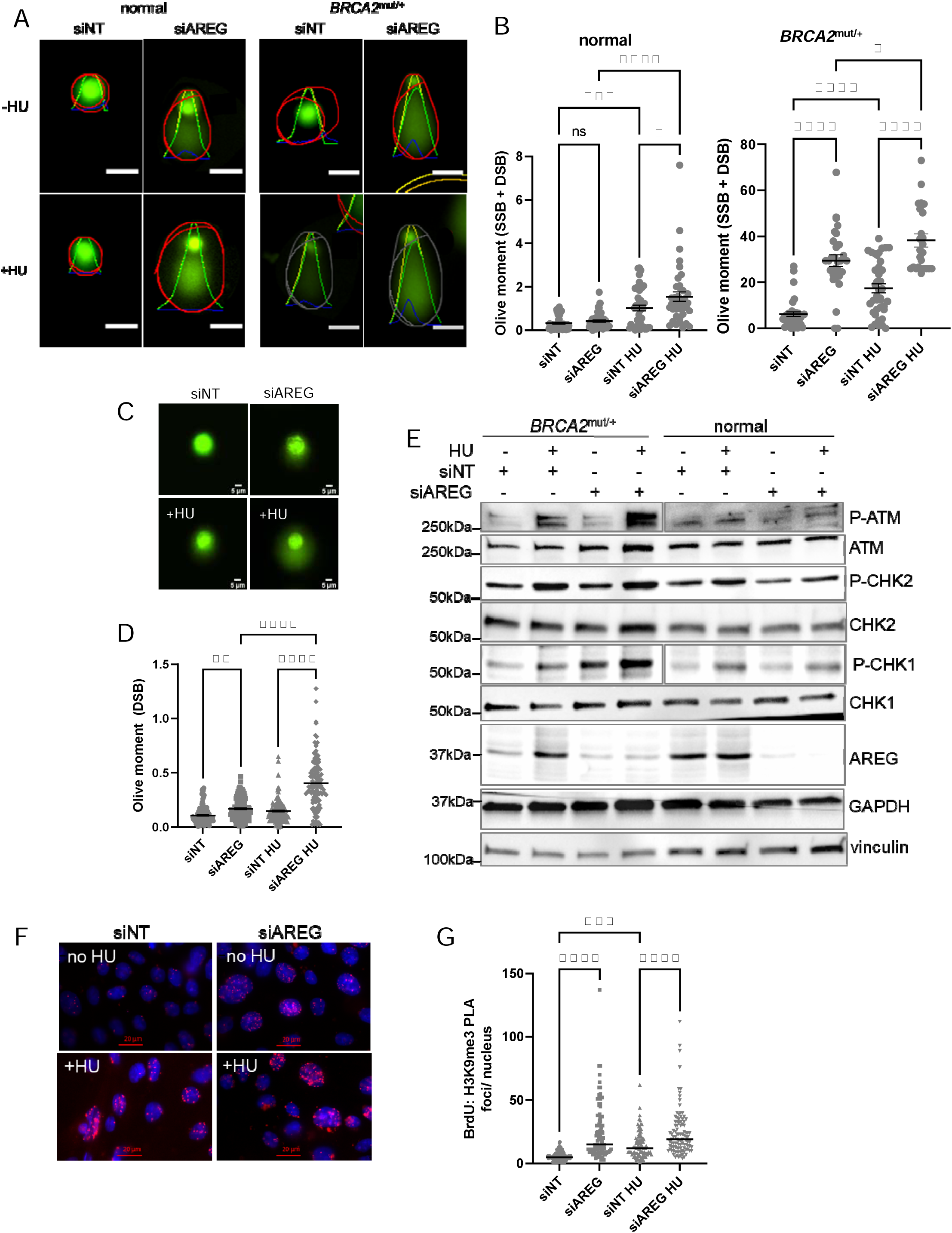
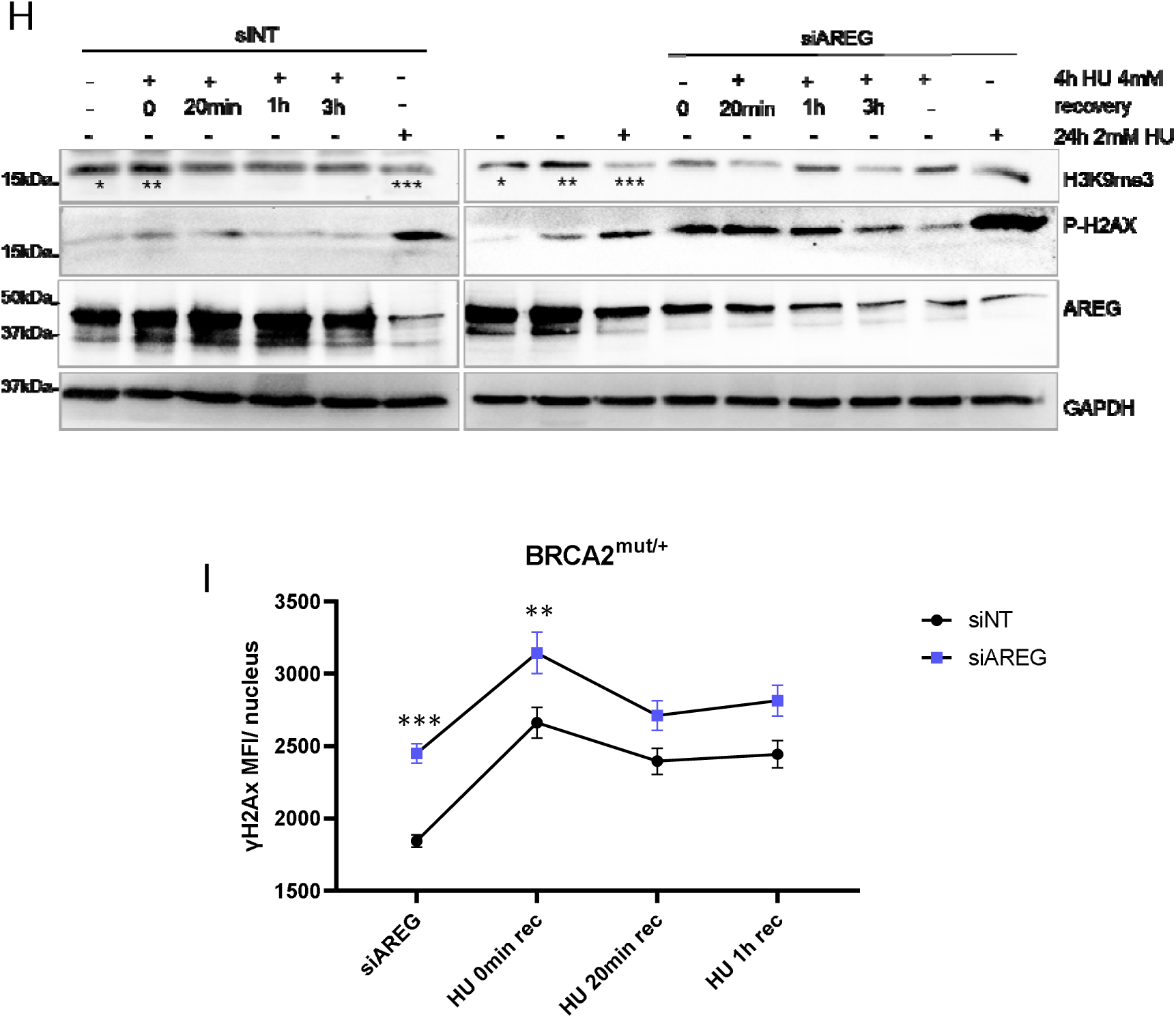
Depletion of AREG promotes genomic instability with or without exogenous replication stress but does not affect H3K9me3 formation at stalled forks or kinetics of DSB repair. **A**, Representative images of comet tails from 3 independent experiments of nuclei within the indicated treatment groups for normal and *BRCA2*^mut/+^ MECs. *BRCA2*^mut/+^ and normal MECs were transfected with siNT or siAREG and 72h later treated with vehicle or 4mM HU (4h) and subjected to an alkaline comet assay to detect ssDNA and dsDNA breaks. Bars=30 pixels. **B,** Graphs depict tail moments (n=50 per group) calculated using the OpenComet plugin in Fiji (https://cometbio.org/). Examples in A show parameters of tail moment calculations based on mean intensity of DNA at the head vs comet tail (green curve), total DNA intensity (red outline) and the intensity of tail DNA (blue curve). *p<.05; **p<.01; ***p<.001; ****p<.0001 One-way ANOVA. **C,** Representative images of nuclei from *BRCA2*^mut/+^ MECs transfected as indicated and electrophoresed nuclei in neutral conditions. **D,** Graph of tail moment derived from neutral comet assays representing DSBs in BRCA2^mut/+^ nuclei treated and analyzed as in A. **p<.01; ****p<.0001; One-way ANOVA. **E,** Representative immunoblot analysis of DNA damage response proteins in siNT or siAREG transfected MECs treated with vehicle or 4mM HU for 4hrs. GAPDH and vinculin proteins were used as loading controls for low and high percentage PAGE. (n=2 independent experiments). **F,** Representative images derived from 2 independent experiments of nuclei from a PLA assay of H3K9me3:IdU-labelled replication forks in *BRCA2*^mut/+^ MECs transfected as indicated and treated with HU for 2h. DNA was labeled with IdU for the final 20 minutes of HU treatment and a kit used for proximity ligation reactions conducted after incubation with anti-H3K9me3 and anti-BrdU according to the manufacturer’s instructions as detailed in Methods. **G,** Graph depicting PLA foci number per nucleus enumerated using Fiji. ***p<.001; ****p<.0001 One-way ANOVA. **H,** Immunoblot of *BRCA2*^mut/+^ MECs transfected with siNT or siAREG then treated as indicated and lysates reacted with anti-H3K9me3, anti-P-H2AX and anti-AREG. GAPDH served as a protein loading control. Lysates from lanes 1*, 2** and 6*** from siNT-transfected MECs were reloaded with siAREG-transfected lysates for direct comparison. **I,** Graph of P-H2AX MFI from IF analysis of each treatment. **p<.01; ***p<.001 One-way ANOVA.

In addition to ssDNA gaps, BRCA2 deficiency results in suboptimal fork protection resulting in DSBs due to resection of ssDNA at stalled forks and fork collapse (38). Both lesions can ultimately convert into DSBs. To assess the effect of AREG depletion on DSBs in *BRCA2*^mut/+^ MECs we used a neutral comet assay. Relative to siNT-transfected MECs, a significant increase in DSBs was induced in AREG-depleted *BRCA2*^mut/+^ MECs in both untreated and in HU-treated conditions (Figures 4C, 4D). Thus, AREG is an important mediator of genomic stability, most prominently in *BRCA2*^mut/+^ MECs where it is critical for preventing excessive DNA damage in response to acute RS.

We next analyzed the activation of DNA damage response proteins. A low but reproducible increase in P-ser1981 ATM (Figure 4E and Supplementary Figure S4) was observed in AREG-depleted *BRCA2*^mut/+^ cells without HU treatment, commensurate with detection of DSBs breaks. An increase in P-CHK1 in *BRCA2*^mut/+^ siAREG-transfected MECs relative to non-targeting siRNA-transfected MECs was also evident in the absence of HU likely consistent with increased ssDNA gaps and RS in these cells. P-ATM, P-CHK2 and P-CHK1 were all activated following HU in siNT-transfected *BRCA2*^mut/+^ MECs and this activation was strongly increased in AREG-depleted MECs (Figure 4E) commensurate with the increased levels of both SSBs and DSBs we observed in comet assays. As expected, HU also induced an increase in the 37kDa and 18kDa AREG in *BRCA2*^mut/+^ MECs. All responses were reduced in normal MECs and immediately following HU although AREG protein increased in normal MECs 3hr post-HU treatment (Figure 1F).

Aside from constitutive heterochromatin, H3K9me3 also forms locally and transiently at stressed replication forks. This helps to maintain fork stability associated with the compaction of chromatin, forming an environment that retains fork protection and remodeling factors until the stress is resolved (34). Given the effect of nuclear-translocated AREG KD on global heterochromatin, we reasoned that AREG KD might impair the transient formation of H3K9me3 specifically at stalled replication forks resulting in increased fork collapse. To investigate this, we performed a proximity ligation assay in *BRCA2*^mut/+^ MECs 72hrs after siAREG or siNT transfection treated with HU for 2h. Nascent DNA was labeled with iodo-deoxyuridine (IdU) followed by incubation with mouse anti-BrdU that detects IdU and rabbit anti-H3K9me3 and colocalization detected with secondary oligo-coupled antibodies then visualized by rolling circle amplification with Alexa564-labelled nucleotides. While untreated/siNT transfected *BRCA2*^mut/+^ MECs exhibit a low but detectable level of BrdU:H3K9me3 proximal foci, as anticipated, HU treatment resulted in a significant increase in focus formation (Figures 4F, 4G). AREG KD alone also increased foci above control levels consistent with RS. Compared to control HU-treated MECs, significantly greater foci numbers were detected in AREG KD MECs following HU treatment. Overall, we conclude that, although transient local formation of heterochromatin at stalled forks is not compromised in AREG-depleted *BRCA2*^mut/+^ MECs, these cells experience increased RS and DSBs.

DSB repair requires H3K9me3 formation around DSBs involves SUV39h1 and HP1α in order to recruit repair complexes and activate ATM (35). To determine if DSB repair kinetics are altered in AREG-depleted MECs with reduced and decompacted heterochromatin, we treated MECs with 4mM HU for 4 hrs followed by a recovery period. Consistent with an increase in DSBs, AREG KD in *BRCA2*^mut/+^ MECs significantly increased P-H2AX compared to siNT-transfected MECs both with and without HU treatment (Figures 4H and 4I and Supplementary Figures S5A, S5B). AREG protein also increased following a 24hr 2mM treatment of MCF-10A spontaneously immortalized MECs while AREG depletion increased P-H2AX compared with untreated cells (Supplementary Figure S5C). The kinetics of the decrease in P-H2AX followed a similar time course in siNT-transfected MECs *BRCA2*^mut/+^ MECs post-HU treatment, however P-H2AX levels persisted at a higher level in AREG-depleted MECs. The higher baseline level of DNA damage in AREG KD MECs is ostensibly due to recurrent DNA breaks as a consequence of dysregulated replication and transcription.

Interestingly, pan-nuclear P-H2AX (homogenous P-H2AX throughout the nucleus) was significantly greater in AREG-depleted, HU-treated MECs (Supplementary Figures S5D, S5E). Pan-nuclear P-H2AX can be mediated by ATM, DNA-PKc or JNK in S phase (67) and be indicative of early stage apoptosis (68) or large scale changes in chromatin structure (69). While apoptotic cells were rare, our results show clearly that AREG KD alters chromatin structure. Thus, the increased nAREG and H3K9me3 in response to RS may be important for avoiding further stress by transiently repressing transcription and slowing replication thus limiting transcription:replication complex collisions. Nevertheless, despite the increase in HU-induced DNA damage associated with a global reduction of H3K9me3, these data are consistent with the ability of *BRCA2*^mut/+^ MECs depleted of AREG to maintain both H3K9me3-dependent stalled fork stabilization and DSB repair activity.

### Endogenous AREG depletion increases new replication origins and firing of multiple origins

Although transient H3K9me3 stalled fork protection appeared unaffected by depletion of AREG, the loss of H3K9me3 and decompaction of heterochromatin could have significant effects on replication and transcription. Therefore, we next used DNA fiber assays to investigate the impact of AREG depletion and associated alteration in heterochromatin on replication in *BRCA2*^mut/+^ MECs. Cells transfected with siNT or siAREG were incubated sequentially with the thymidine analogues CldU followed by IdU for 20 minutes each. A small but significant decrease in ldU:CldU ratio was observed after AREG KD which may be consistent with exacerbation of fork reversal which occurs when BRCA2 is depleted (70). To address the effect of AREG depletion on strand resection, cultures were treated with HU for 4hr immediately after IdU washout. Strand resection occurred in both siNT and siAREG transfected MECs, however this was significantly greater in siAREG-depleted MECs (Figures 5A, 5B). Fork regression was prevented by the co-treatment with the MRE11 inhibitor, mirin, although protection was incomplete in siAREG-transfected MECs, possibly attributed to an increased frequency of fork collapse in the AREG KD MECs. Notably, DNA fiber lengths were significantly shorter in untreated siAREG-transfected MECs compared with control MECs indicating a slowed replication rate (Figure 5C). Slowing of the replication fork can result in activation of normally dormant replication origins (71). Collapsed forks resulting in restart at new origins as indicated by IdU-only labeled fibers were also significantly more frequent in AREG-depleted (Figure 5D). No changes between treatment groups in bidirectional forks were observed (not shown), however AREG KD resulted in a significantly greater number of fibers with CldU-IdU-CldU-IdU-labeling indicative of multiple origins of replication within a single fiber (Figures 5E, 5F).

**Figure 5.**
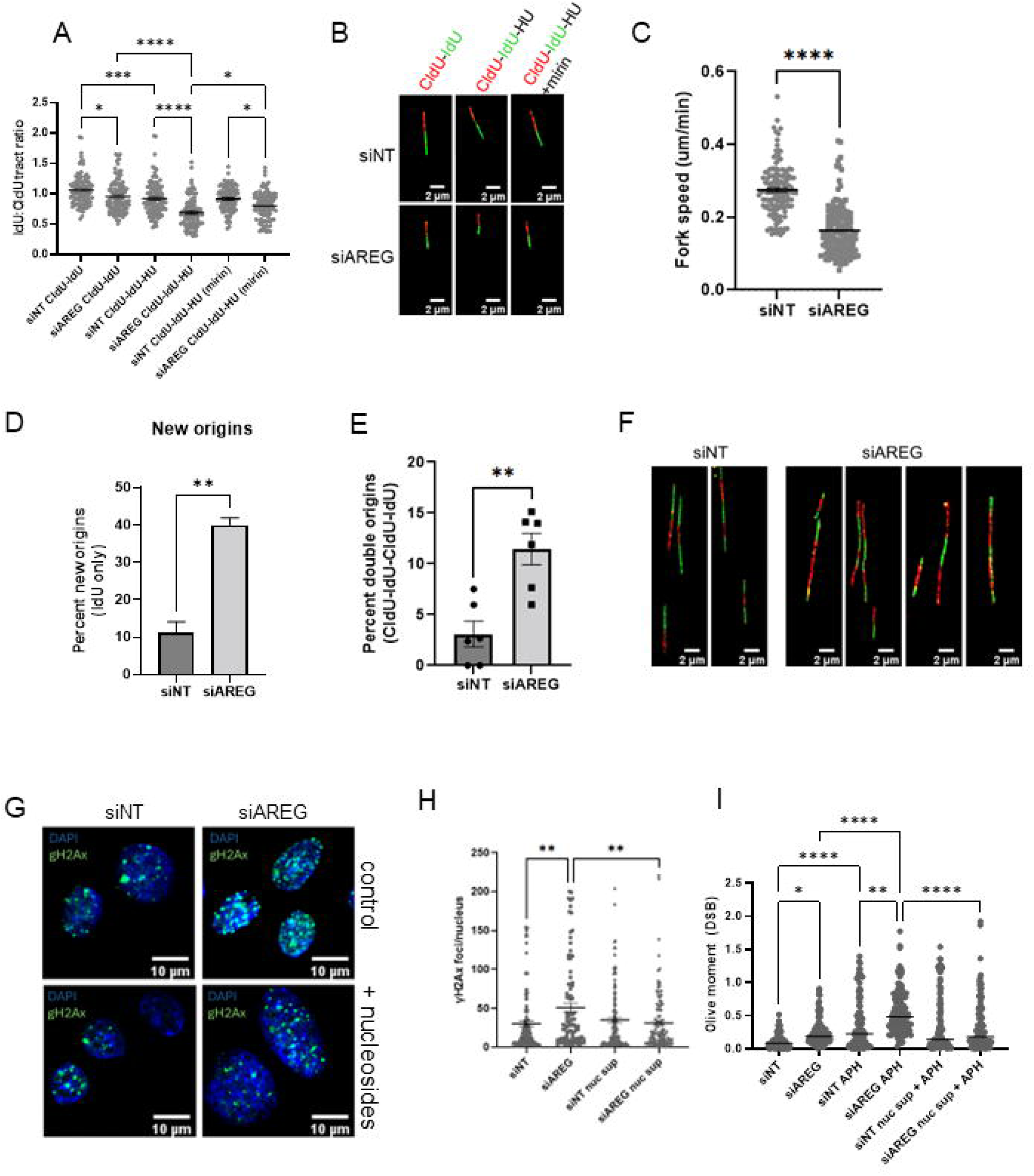
Loss of endogenous AREG depletes nucleotides reducing fork stability and slowing replication fork progression. DNA fiber assays were performed on *BRCA2*^mut/+^ MECs transfected with siNT or siAREG. Cells were incubated sequentially with the thymidine analogues CldU followed by IdU for 20 minutes in the presence of exogenous AREG. Some cultures were treated with HU for 4h immediately after IdU washout. **A,** CldU (red) and IdU (green) tract lengths were measured and the IdU:CldU ratio calculated (100 tracts per group). Relative to siNT transfected MECs, siAREG-transfection resulted in a significant decrease in tract length after HU treatment indicating nucleolytic degradation of ssDNA. Fork degradation was prevented by the co-treatment with the MRE11 inhibitor mirin in control MECs but only partially inhibited in AREG-depleted MECs suggesting fork collapse. *p<.05; ***p<.001; ****p<.0001 One-way ANOVA. **B,** Representative images of fibers from cells treated as indicated. Bars=2µm. **C,** Fiber lengths were measured and used to calculate replication speed (CldU tract+IdU tract)/2/20min=µm/min) in siAREG and siNT-transfected MECs. ****p<.0001 two-tailed t-test. **D,** Graph shows percentage of new origins indicative of collapsed forks labeled with IdU (green) only (yellow arrow) as indicated in the adjacent image. Each data point is the mean percentage of 20 tracts derived from 5 random microscopic fields. *p< .05; two-tailed t-test. **E,** Graph showing the percentage of tracts with multiple origins of replication within a single strand labeled with CldU-IdU-CldU-IdU Each data point represents the mean of 20 tracts counted from random microscopic fields from 3 independent experiments (n=120). **p<.01; two-tailed t-test. **F,** Representative images of bidirectional forks and tracts with multiple origins of replication in siNT and siAREG transfected MECs (blue arrows) Bars=2µm. **G,** Representative images (n=3 independent experiments) of P-H2AX foci in MECs transfected with siNT or siAREG in the absence (control) or presence of supplemental nucleoside. Bar=10µm. **H,** Graph of P-H2AX foci in a minimum of 120 cells per control group and AREG-depleted *BRCA2*^mut/+^ MECs cultured in the presence or absence of supplemental nucleoside. **p<.01; ****p<.0001 One-way ANOVA. **I,** Graph of neutral comet assay tail moments from *BRCA2*^mut/+^ MECs treated for 20h with 0.4 µM APH to induce RS in the presence or absence of nucleoside supplementation to the media. n= minimum 120 nuclei per group. *p<.05; **p<.01; ****p<.0001 One-way ANOVA.

The genome is sequentially replicated with open chromatin replicating first followed by heterochromatic regions in late S phase (72). Dysregulated replication across the genome could result in nucleotide depletion thereby promoting RS and DNA damage. In support of this, supplementation of media with nucleosides significantly reduced P-H2AX foci accumulation following AREG KD (Figures 5G, 5H). Using a neutral comet assay, we also tested the effect of nucleotide supplementation directly on DSB formation following RS induced by a 24hr treatment with the DNA polymerase α inhibitor, aphidicolin. Figure 5I indicates a significant increase in DSBs in siAREG-transfected *BRCA2*^mut/+^ MECs compared to normal MECs. Similar to HU-induced RS, 20hrs of culture in the presence of aphidicolin increased DSBs significantly more in AREG KD compared to control MECs. Again, consistent with a role for nucleotide depletion in these cells, nucleoside supplementation for 48 hrs markedly reduced the Olive tail moment in AREG KD but had no effect on siNT-transfected MECs.

Reduced constitutive heterochromatin is expected to increase global transcription levels which in turn can result in replisome:transcription complex collisions and R-loop formation. Indeed, increased global transcription can result in replication stress (73). Transfection of cells with AREGΔC11 was previously shown to reduce global transcription (9). To determine if RNA transcription was increased by AREG depletion, we cultured siNT and siAREG-transfected MECs in the presence of BrU for 20 min followed by IF detection with anti-BrdU. IF analysis in Fiji (Supplementary Figures S5A and S6B, respectively) show that the level of transcription increased significantly in AREG-depleted MECs compared to controls.

In total, these results are consistent with reduction and decompaction of H3K9me3 heterochromatin resulting in dysregulation of replication, enhanced RS and increased genomic instability in AREG-depleted MECs.

### Absence of AREG in BRCA2^mut/+^ MECs disrupts the Ran-GTPase gradient, reduces prelamin A maturation and promotes extensive nucleoplasmic reticulum

The association of AREG with lamin A and reduction of H3K9me3 and heterochromatin compaction after AREG depletion suggested that the effects of depletion might impact the NM itself. Heterochromatin was found to be essential for maintaining the Ran gradient and the nuclear import of ATM in fibroblasts from Hutchinson-Gilford progeria syndrome patients (74). The ratio of nuclear to cytoplasmic Ran is typically 3:1. Ran IF in siNT-transfected control MECs in AREG KD MECs revealed the normal nuclear-cytoplasmic distribution. However, a profound reduction of the Ran gradient was observed in AREG KD *BRCA2*^mut/+^ (Figures 6A, 6B).

**Figure 6.**
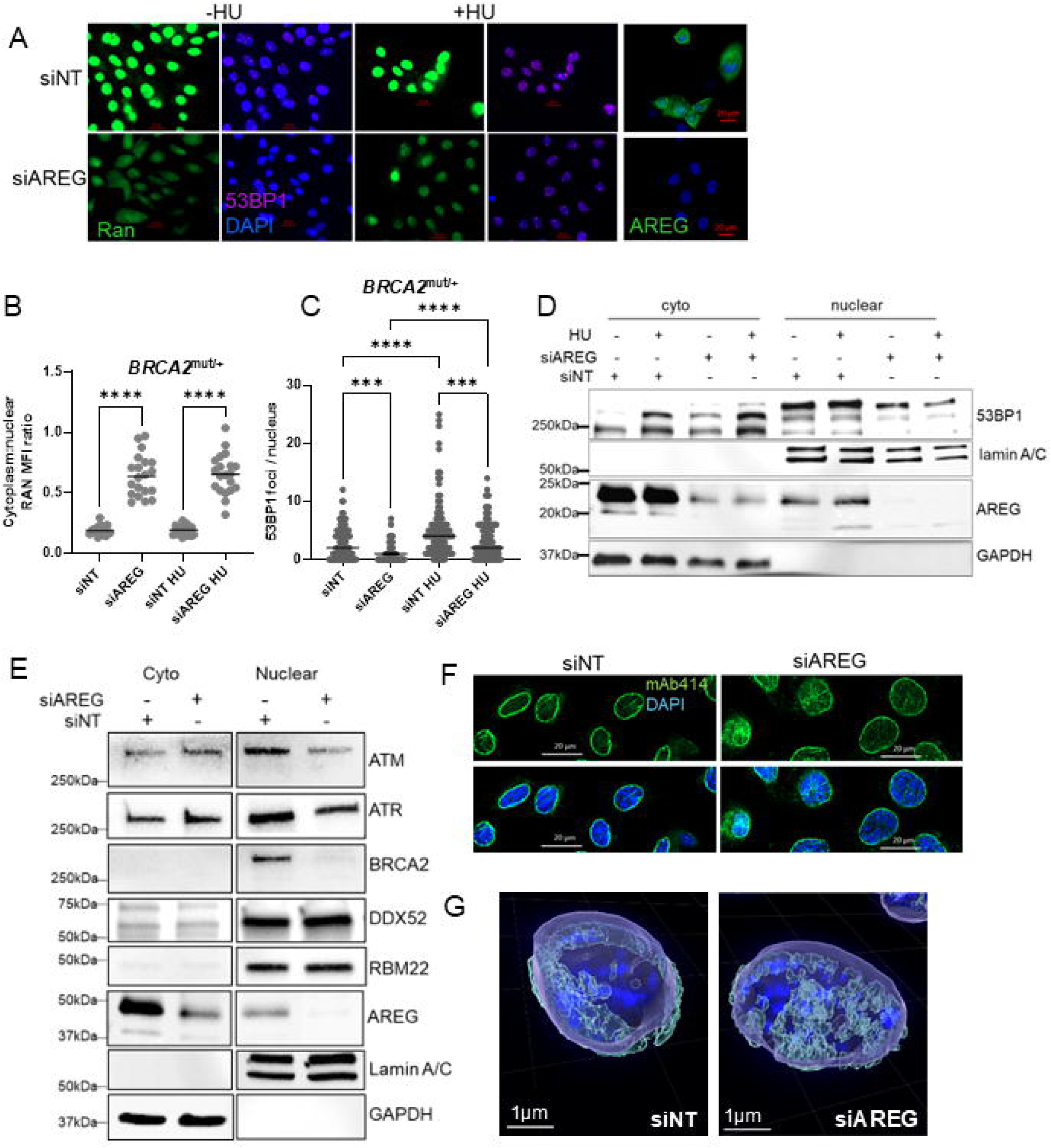
AREG depletion dissipates the nuclear Ran gradient and promotes nuclear membrane invagination. **A**, IF using antibodies to Ran and 53BP1 in *BRCA2*^mut/+^ MECs 72h after transfection with siNT or siAREG. **B,** Graph of mean cytoplasmic to nuclear ratio fluorescence intensity of Ran in MECs transfected as indicated MECs (n=20 cells per treatment group) representative of 3 independent experiments. **C,** Representative graph depicting 53BP1 foci per nucleus in MECs transfected as indicated and treated with vehicle or 4mM HU for 4h prior to fixation. (n=100 nuclei from 3 independent experiments). B and C; ***p<.001; ****p<.0001 (One-way ANOVA) **D,** Immunoblot of cytoplasmic and nuclear fractions from *BRCA2*^mut/+^ MECs transfected with siNT or siAREG treated +/-HU probed with anti-53BP1, AREG, lamin A/C and GAPDH. AREG-HE is a high exposure of the lower ∼18kDa AREG band in the nuclear fraction. **E,** Immunoblots for the indicated proteins in cytoplasmic and nuclear fractions from siNT and siAREG transfected MECs. Note that high mol wt ATR, ATM and BRCA2 are reduced in nuclei following AREG KD. The AREG peptide band shown is the 23kDa species. **F,** IF of FG nucleoporins (mAb414) in MECs transfected and treated with HU as indicated. **G,** Imaris rendering of confocal images of siNT and siAREG-transfected MECs immunostained with mAb414 to detect FG nucleoporins (green) counterstained with DAPI (blue).

53BP1 helps mediate ATR-Chk1-p53 signaling to protect replication forks in response to RS (75). Nuclear localization of 53BP1 is dependent on active transport mediated by Ran-GTPase-importin α/β-mediated nuclear import (76). Consistent with their BRCA2-deficient status, we found that unperturbed *BRCA2*^mut/+^ MECs also contained nuclear 53BP1 foci (Figures 6A). Although, loss of endogenous AREG significantly decreased 53BP1 foci (Figure 6A, 6C). Given the disruption of the Ran gradient, we next investigated whether reduced nuclear foci might be a consequence of altered 53BP1 subcellular localization. Immunoblot of cytoplasmic and nuclear fractions of control and AREG-depleted *BRCA2*^mut/+^ MECs revealed cytoplasmic 53BP1 in both siNT and siAREG transfected MECs (Figure 6D). 53BP1 is SUMOylated in response to DNA damage (77) and high molecular weight forms were increased in response to either AREG KD and/or HU treatment. However, nuclear 53BP1 protein levels were consistently lower in AREG-depleted MECs regardless of HU-induced RS. Note that, as in Figure 1, the 23kDa and 18kDa nAREG peptides were increased by HU treatment.

ATM and ATR proteins also require Ran-GTPase facilitated nuclear transport through the nuclear pore. Although we cannot rule out that a portion of the P-ATM induced by HU-treatment following AREG depletion (Figure 4E) was cytoplasmic, the nuclear content of both ATM and ATR was reduced (Figure 6E). We did not detect BRCA2 (Figure 6D) in the nuclear or cytoplasmic fractions of these MECs. In contrast, AREG KD did not impact the nuclear content of the small nuclear proteins RBM22 (pre-mRNA splicing) and DDX52 (RNA helicase). Thus, in addition to replication dysregulation in AREG-depleted MECs, the reduced Ran gradient and import of DNA repair proteins could also negatively impact on genome repair capacity.

Heterochromatin interactions with the nuclear lamina and nuclear pore complexes maintain its peripheral localization (78) and also affects the structural integrity of the nuclear envelope which, when compromised, results in increased NM invagination (79) and rupture (80). Since KD of AREG reduces compaction and levels of H3K9me3 heterochromatin and compromises the Ran gradient and nuclear import in AREG-depleted MECs, we next looked to see if NPC distribution was altered in these cells. The images in Figure 6F show the primarily uniform nuclear rim staining in control-transfected MECs with mAb414 which recognizes FG repeat nucleoporins (Nups). AREG depletion resulted in a reduction in Nup IF at the nuclear rim and introduced regions of discontinuity. In addition, these cells demonstrated increased nucleoplasmic Nup distribution which has been attributed to NM invaginations also known as nucleoplasmic reticulum (NR) (81). Blockage of lamin farnesylation (which reduces lamin A incorporation into the nuclear matrix (82)) can increase the formation of NR (81) as does inhibition of the ZMPSTE24 endoprotease increasing farnesylated prelamin A (83). 3-D images in Imaris from z-stacks of MECs immunostained with mAb414 following siNT or siAREG transfection shows a typical cell following AREG depletion showing enhanced formation of NR (Figure 6G). In a subset of transfections NR invaginations were large enough to be detected within z-stacks of lamin A/C IF (Supplementary Figure S7).

### AREG deficiency activates IFN-1-like gene expression and increases senescence following RS

Loss of heterochromatin domains can result in NM blebbing and ruptures (80). Evaluation of nuclear blebs and ruptures following AREG KD in *BRCA2*^mut/+^ MECs showed an increased incidence compared to control that was not further enhanced upon acute HU-induced RS (Figures 7A, 7B). Notably, we also detected cGAS accumulation at DAPI stained cytoplasmic DNA which has potential to activate an IFN-like response. Although infrequent, anaphase bridges occurred in AREG-depleted but not control MECs (panel v) consistent with RS and incomplete replication (84).

**Figure 7.**
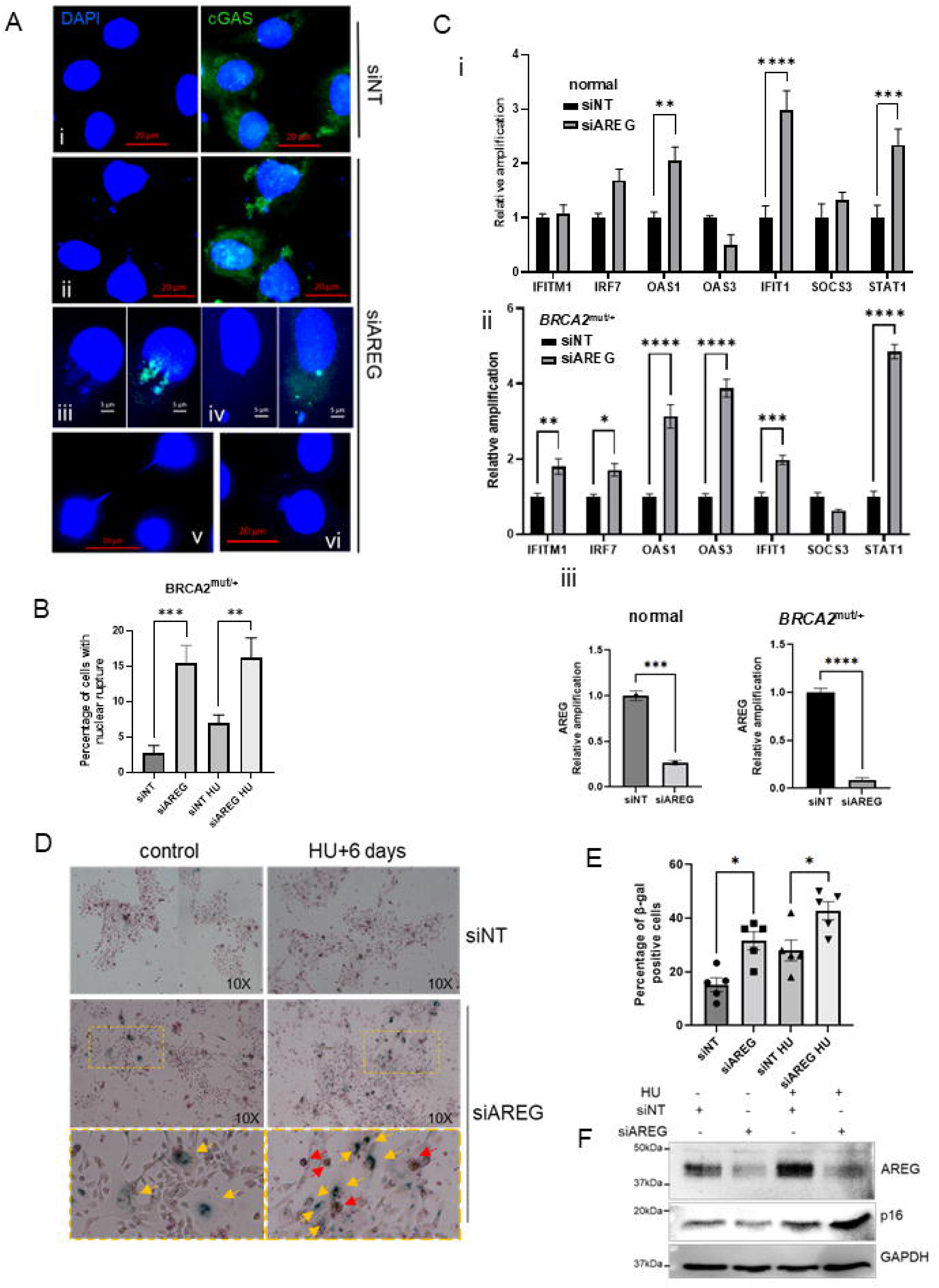
AREG supports nuclear membrane integrity preventing an IFN-1-like response and reduces senescence following replication stress. **A**, Representative images of DAPI and cGAS immunostained nuclei from (i), siNT transfected MECs and (ii-v), siAREG-transfected MECS. (vi) AREG depletion also resulted in the appearance of anaphase bridges not detected in controls. **B,** Graph of percentage of MECs with nuclear ruptures after siNT or siAREG transfection treated with or without HU. Percentages are derived from minimum of 233 nuclei per group and each data point represents the mean percentage of nuclei with ruptures calculated from a minimum of 3 microscopic fields compiled from 3 independent experiments. Bars are S.E.M. ***p<.001; **p<.01 One-way ANOVA. **C,** (i) qPCR for IFN-like pathway genes in *BRCA2*^mut/+^ MECs and (ii) normal MECs 72hrs following siNT and siAREG transfection. **p<.01 ***p<.001; ****p<.0001; One-way ANOVA. Bars are S.E.M. (triplicates from 2 independent experiments) (ii) Graph of AREG qPCR following transfections. D, *BRCA2*^mut/+^ MECs were transfected as shown and treated with 4mM HU for 4 hrs. Media was changed and cells stained to detect SAβ-gal activity 4 days later. Arrows indicate SAβ-gal activity-positive MECs with typical senescent cell morphology. **E,** Quantification of SAβ-gal positive MECs. Each data point represent the average of 50 SAβ-gal-positive MECs (n=250 cells per group). Bars are S.E.M. * p<.05 One-way ANOVA. **E,** RNA from *BRCA2*^mut/+^ MECs treated with vehicle or 15µM HU for 4 days was used for qPCR to detect AREG mRNA. Bars are S.E.M of 3 independent experiments. ****p<.0001; two-tailed t-test. **F,** Immunoblot of AREG and p16 treated as in **D**. GAPDH detection was used as a protein loading control.

Chronic unrepaired DNA damage can induce RS can lead to accumulation of cytosolic DNA which activates the cGAS/STING cytosolic DNA sensing pathway and IFN-1-like pathway gene expression (85), implicated in both neoplasia (86) and senescence (87). Moreover, RNA-DNA hybrids aberrantly accumulate in the cytoplasm after R-loop processing induce the innate immune response (88). H3K9 methytransferase-deficient MEFs display activation of IFN signaling, including overexpression of type I IFN regulators Irf7, Irf9, and Stat1 (89). Given the reduction and/or decompaction of H3K9me3 after AREG KD, NM ruptures and cytosolic DNA in MECs depleted of AREG we analyzed the expression of several genes in the IFN-1 pathway 72 hrs after siAREG or siNT transfection. qRT-PCR revealed AREG-depletion in normal MECs resulted in increased IFIT1 mRNA transcripts in particular as well as STAT1 (Figure 7C(i)). Similarly, AREG-depletion in *BRCA2*^mut/+^ MECs increased IFIT1 and STAT1 mRNAs although relative induction of STAT1 was greater than in normal MECs while OAS1/3 mRNAs were differentially induced in *BRCA2*^mut/+^ MECs (Figure 7C (ii)). qRT-PCR confirmed AREG KD in normal and *BRCA2*^mut/+^ MECs (Figure 7C (iii)).

cGAS activity contributes to senescence establishment (87) and persistent DNA damage as a consequence of RS could ultimately promote senescence. To investigate a role for AREG in protecting *BRCA2*^mut/+^ MECs from RS-induced DNA damage and senescence, we transfected *BRCA2*^mut/+^ MECs with siAREG for 72 hrs then treated with 4mM HU for 4hrs to induce RS. Cell lysates collected 3 days after HU for analysis of the early senescence marker, p16. Figure 7D shows that, after 3 days, siAREG-transfected MECs and siNT controls expressed similar low levels of p16. Exposure to HU induced a small increase in p16 in control cells but prominent induction of p16 in AREG-depleted MECs was evident indicating the onset of senesence. MECs were stained to detect senescence-associated (SA)β-gal activity which was almost undetectable in untreated MECs. Although evidence of (SA)β-gal+ cells showing typical flattened morphology and vesicle formation was present in some siNT-transfected MECs, a significant increase was detected in siAREG-expressing cultures which was further increased in MECs that had been treated with HU. Thus, AREG is critical for constitutive maintenance of genomic integrity in MECs, and prevention of excessive DNA damage following RS which can result in senescence and of key importance in *BRCA2*^mut/+^ MECs sensitive to RS.

## Discussion

This study demonstrates that AREG is essential for maintaining heterochromatin in MECs to protect these cells from genomic instability and maintain NM function. Heterochromatin has critical roles in epigenomic inheritance, maintenance of differentiation and genomic stability (90) at least in part by regulating replication timing (13). The maintenance of H3K9me3 involves the recruitment of SUV39h1 to parental histones on newly synthesized DNA and binding of HP1α which recruits additional histone modifying enzymes (21) and undergoes multimerization (21,22) to bridge and stabilize heterochromatic domains, 24). We discovered that loss of AREG in MECs reduces SUV39h1 and HP1α resulting in heterochromatin loss and decompaction. Thus, during successive replicative cycles, the depletion of these factors in AREG KD MECs is predicted to prevent the normal maintenance of heterochromatin, predisposing these cells to RS which is enhanced in *BRCA2*^mut/+^ MECs. Altered replication dynamics and nucleotide depletion provided functional evidence of heterochromatin loss including reduced fork speed, increased new and multiple origins and ultimately fork stalling and collapse resulting in DSBs in AREG KD MECs.

The stress kinase p38α provides an important rapid signal in response to DNA damage and we have established that the RS-induced increase in AREG colocalization with early endosomes, AREG processing and H3K9me3 formation are p38 activity-dependent. Following phorbol ester stimulation, AREG undergoes ER retrieval to the trans-Golgi network (TGN) and enters the nucleus via nuclear pores based given the presence of an NLS (9). Our data show the colocalization of AREG and lamin A in cytoplasmic and nuclear PLA foci after HU suggesting that, post endocytosis, AREG and prelamin A engage in a common trafficking pathway. Interestingly, sorting nexin 6 (SNX6) is a component of the retromer which is involved in transporting transmembrane receptors from endosomes to the TGN (28). Prelamin A nuclear import and nuclear envelope incorporation are regulated by SNX6 linking it to the outer surface of the ER and is required for lamin A nuclear import through nuclear pores (27). Based on this and the present data, it is plausible that DNA damage-induced retrograde AREG transport facilitates interaction between AREG and prelamin A:SNX6-retromers formed on the cytoplasmic side of the ER membrane, thereby leading to nuclear entry of both proteins. Although HU results in a small increase in AREG mRNA, the trafficking of AREG via this pathway would contribute to rapid stabilization of existing AREG protein during RS due to bypass of the late endosome/lysosomal pathway.

Prelamin A farnesylation, carboxymethylation and cleavage can occur either at the outer surface of the ER or in the nucleus at the nuclear rim (91). Since we found that prelamin A and AREG are both markedly increased in the NM after RS, cleavage of prelamin A may be transiently reduced by the increased association and trafficking with AREG. In the absence of AREG, we observed prelamin A in the cytoplasm which likely represents accumulation of ER-associated prelamin A unable to traffic to the nucleus. Further studies are necessary to resolve these possibilities.

In addition to maintenance, the formation of transient heterochromatin is also an acute response to both DSBs (35) as well as stalled replication forks (34). Although depletion of AREG reduces global heterochromatin and/or compaction, transient heterochromatin formation at stalled forks appeared unaffected based on proximity ligation analysis of H3K9me3 coupled to IdU-labeled nascent DNA. Resolution of P-H2AX occurs in AREG-depleted MECs with similar kinetics as controls following withdrawal of HU. This suggests that sufficient levels of HP1α and SUV39h1 remain to allow for a portion of DSB repair in euchromatin however the higher basal level of P-H2AX is likely due to the de novo rate of DSB formation in AREG KD MECs.

PLA to detect lamin A:AREG proximity indicated a major cytoplasmic and nuclear corecruitment of AREG and lamin A in response to RS where foci accumulated outside the nucleus, at the periphery and interior. Progerin, non-farnesylated and farnesylated prelamin A have been shown to bind HP1α (30,90) and can stabilize the protein resulting in increased H3K9me3 formation (30). HP1α binding to SUV39h1 also promotes its stabilization (56,57) which may account for the reduction in this methyltransferase in AREG-depleted MECs. Lamin A mutants that fail to incorporate into the nuclear lamina have been shown to destabilize HP1α (93) which is compatible with the nuclear lamina incorporation of prelamin A and AREG in response to RS and reduction in prelamin A delivery to the nucleus in the absence of AREG. Notably HP1α was also reported to promote efficient homologous recombination (94), which is consistent with a nAREG-associated mechanism preventing excessive DNA damage after RS in *BRCA2*^mut/+^ MECs.

The positioning of heterochromatin at the nuclear periphery occurs, in part, in a lamin A/C-dependent manner mediated by PRR14 which interacts with H3K9me3 via HP1α (16). Given this, reduced HP1α would also contribute to both the reduction of peripheral heterochromatin and decondensation we observed in AREG-depleted MECs.

Loss of heterochromatin has been shown to result in dissipation of the Ran gradient and affect the ability to import numerous proteins including including 53BP1 and ATM (74) and would indeed impair lamin A import (27). Thus, the compromised Ran gradient could reduce the repair capacity in MECs contributing to increased DNA breaks, nevertheless, despite the reduction in nuclear ATM and reduction in 53BP1 foci in AREG KD MECs, activation of ATM and H2AX foci were evident following AREG KD and markedly increased following HU treatment in *BRCA2*^mut/+^ MECs.

NM rupture releasing DNA into the cytoplasmic compartment, likely combined with RS generated cytoplasmic DNA fragments in AREG KD MECs, resulted in robust activation of the IFN-1-like response and development of SA-βgal enzymatic activity. The cGAS/STING pathway is involved in enforcing senescence (87). Thus, by both maintaining and increasing heterochromatin in response to RS, nAREG prevents excessive DNA damage and senescence. Indeed, this feature of nAREG could also contribute to chemoresistance. Similar to induction of AREG mRNA by HU in *BRCA2*^mut/+^ MECs, radiation resistant pancreatic cancer cells were shown to express approximately 2.5X the level of AREG mRNA compared to sensitive cells (95). AREG upregulation was also shown in cisplatin-resistant MCF-7 cells (96). Moreover, nuclear pro-AREG has specifically been shown to mediate chemoresistance in pancreatic cancer cells (97).

During differentiation, H3K9me3 heterochromatin forms large “islands” in the genome in a cell-type-specific manner (98). One function of these heterochromatic regions is to form a barrier to transcription-factor-mediated cell-type reprogramming and in doing so maintain the cell-type identity of differentiated cells (90). Although mature luminal MECs are fully differentiated, they express the highest level of AREG compared with progenitor cells (97,98). Thus, an intriguing possibility is that, as progenitors differentiate, increasing endogenous AREG expression could help to maintain lineage fidelity.

Repeated exposure to RS is thought to contribute significantly to the development of cancer as a consequence of ensuing genomic instability (31,101). Given this, our study indicates that nAREG is a critical factor that protects both normal and especially *BRCA2^m^*^ut/+^ MECs from DNA damage that could result in senescence and could potentially be used as a marker of RS.

## Acknowledgements

This work was funded by Canadian Cancer Society Research Institute grants i2i-18 #705975 and INNOV-18 #705940 to M.AC.P. We thank the University of Ottawa CBIA Core facility for assistance with imaging and image analysis. We are grateful to Cheryl Lewis (UT Southwestern, Texas) for the generous gift of hTERT-mammary epithelial cells and William Muller (McGill University, Quebec) for MMTV-cre mice.

## Author Contributions

T. J. performed experiments, all image analysis, managed animal husbandry and contributed to experimental design and manuscript writing. A.Z. performed experiments. M.A.C.P. designed the research, experiments and wrote the manuscript. The authors have no competing interests.

## Notes

### Competing Interest Statement

The authors have declared no competing interest.

### Summary of Updates

Figure 8 has been removed and only the IHC for AREG is now presented in Figure 1. Several figures have been reordered. New data and accompanying text (revised manuscript Figure 3K and 3N) more clearly define the role and relationship of HP1 alpha and prelamin A in the AREG-dependent response to replication stress.

## References

1. Singh SS, Chauhan SB, Kumar A, Kumar S, Engwerda CR, Sundar S, et al. Amphiregulin in cellular physiology, health, and disease: Potential use as a biomarker and therapeutic target. Journal Cellular Physiology [Internet]. 2022 Feb [cited 2024 Oct 24];237(2):1143–56. Available from: https://onlinelibrary.wiley.com/doi/10.1002/jcp.30615

2. Ciarloni L, Mallepell S, Brisken C. Amphiregulin is an essential mediator of estrogen receptor alpha function in mammary gland development. Proc Natl Acad Sci U S A. 2007 Mar 27;104(13):5455–60.

3. Booth BW, Boulanger CA, Anderson LH, Jimenez-Rojo L, Brisken C, Smith GH. Amphiregulin mediates self-renewal in an immortal mammary epithelial cell line with stem cell characteristics. Exp Cell Res. 2010 Feb 1;316(3):422–32.

4. Berasain C, Avila MA. Amphiregulin. Semin Cell Dev Biol. 2014 Apr;28:31–41.

5. Fukuda S, Nishida-Fukuda H, Nakayama H, Inoue H, Higashiyama S. Monoubiquitination of pro-amphiregulin regulates its endocytosis and ectodomain shedding. Biochem Biophys Res Commun. 2012 Apr 6;420(2):315–20.

6. Johnson GR, Saeki T, Auersperg N, Gordon AW, Shoyab M, Salomon DS, et al. Response to and expression of amphiregulin by ovarian carcinoma and normal ovarian surface epithelial cells: nuclear localization of endogenous amphiregulin. Biochem Biophys Res Commun. 1991 Oct 31;180(2):481–8.

7. Kimura H. Schwannoma-derived growth factor must be transported into the nucleus to exert its mitogenic activity. Proc Natl Acad Sci U S A. 1993 Mar 15;90(6):2165–9.

8. Olsnes S, Klingenberg O, Więdłocha A. Transport of Exogenous Growth Factors and Cytokines to the Cytosol and to the Nucleus. Physiological Reviews [Internet]. 2003 Jan 1 [cited 2024 Oct 24];83(1):163–82. Available from: https://www.physiology.org/doi/10.1152/physrev.00021.2002

9. Isokane M, Hieda M, Hirakawa S, Shudou M, Nakashiro K, Hashimoto K, et al. Plasma-membrane-anchored growth factor pro-amphiregulin binds A-type lamin and regulates global transcription. J Cell Sci. 2008 Nov 1;121(Pt 21):3608–18.

10. Janssen A, Colmenares SU, Karpen GH. Heterochromatin: Guardian of the Genome. Annu Rev Cell Dev Biol [Internet]. 2018 Oct 6 [cited 2024 Oct 24];34(1):265–88. Available from: https://www.annualreviews.org/doi/10.1146/annurev-cellbio-100617-062653

11. Grewal SIS. The molecular basis of heterochromatin assembly and epigenetic inheritance. Mol Cell. 2023 Jun 1;83(11):1767–85.

12. de Koning APJ, Gu W, Castoe TA, Batzer MA, Pollock DD. Repetitive elements may comprise over two-thirds of the human genome. PLoS Genet. 2011 Dec;7(12):e1002384.

13. Rhind N, Gilbert DM. DNA replication timing. Cold Spring Harb Perspect Biol. 2013 Aug 1;5(8):a010132.

14. Van Steensel B, Belmont AS. Lamina-Associated Domains: Links with Chromosome Architecture, Heterochromatin, and Gene Repression. Cell [Internet]. 2017 May [cited 2024 Oct 24];169(5):780–91. Available from: https://linkinghub.elsevier.com/retrieve/pii/S0092867417304737

15. Briand N, Collas P. Lamina-associated domains: peripheral matters and internal affairs. Genome Biol [Internet]. 2020 Dec [cited 2024 Oct 24];21(1):85. Available from: https://genomebiology.biomedcentral.com/articles/10.1186/s13059-020-02003-5

16. Poleshko A, Mansfield KM, Burlingame CC, Andrake MD, Shah NR, Katz RA. The human protein PRR14 tethers heterochromatin to the nuclear lamina during interphase and mitotic exit. Cell Rep. 2013 Oct 31;5(2):292–301.

17. Solovei I, Wang AS, Thanisch K, Schmidt CS, Krebs S, Zwerger M, et al. LBR and lamin A/C sequentially tether peripheral heterochromatin and inversely regulate differentiation. Cell. 2013 Jan 31;152(3):584–98.

18. Padeken J, Methot SP, Gasser SM. Establishment of H3K9-methylated heterochromatin and its functions in tissue differentiation and maintenance. Nat Rev Mol Cell Biol [Internet]. 2022 Sep [cited 2023 Sep 17];23(9):623–40. Available from: https://www.nature.com/articles/s41580-022-00483-w

19. Alabert C, Groth A. Chromatin replication and epigenome maintenance. Nat Rev Mol Cell Biol. 2012 Feb 23;13(3):153–67.

20. Loyola A, Tagami H, Bonaldi T, Roche D, Quivy JP, Imhof A, et al. The HP1alpha-CAF1-SetDB1-containing complex provides H3K9me1 for Suv39-mediated K9me3 in pericentric heterochromatin. EMBO Rep. 2009 Jul;10(7):769–75.

21. Bannister AJ, Zegerman P, Partridge JF, Miska EA, Thomas JO, Allshire RC, et al. Selective recognition of methylated lysine 9 on histone H3 by the HP1 chromo domain. Nature. 2001 Mar 1;410(6824):120–4.

22. Lachner M, O’Carroll D, Rea S, Mechtler K, Jenuwein T. Methylation of histone H3 lysine 9 creates a binding site for HP1 proteins. Nature. 2001 Mar 1;410(6824):116–20.

23. Canzio D, Chang EY, Shankar S, Kuchenbecker KM, Simon MD, Madhani HD, et al. Chromodomain-mediated oligomerization of HP1 suggests a nucleosome-bridging mechanism for heterochromatin assembly. Mol Cell. 2011 Jan 7;41(1):67–81.

24. Schotta G, Ebert A, Krauss V, Fischer A, Hoffmann J, Rea S, et al. Central role of Drosophila SU(VAR)3-9 in histone H3-K9 methylation and heterochromatic gene silencing. EMBO J. 2002 Mar 1;21(5):1121–31.

25. Dittmer TA, Misteli T. The lamin protein family. Genome Biol. 2011;12(5):222.

26. Naetar N, Ferraioli S, Foisner R. Lamins in the nuclear interior − life outside the lamina. Journal of Cell Science [Internet]. 2017 Jul 1 [cited 2022 Feb 20];130(13):2087–96. Available from: https://journals.biologists.com/jcs/article/130/13/2087/56175/Lamins-in-the-nuclear-interior-life-outside-the

27. González-Granado JM, Navarro-Puche A, Molina-Sanchez P, Blanco-Berrocal M, Viana R, De Mora JF, et al. Sorting Nexin 6 Enhances Lamin A Synthesis and Incorporation into the Nuclear Envelope. Larijani B, editor. PLoS ONE [Internet]. 2014 Dec 23 [cited 2025 Mar 17];9(12):e115571. Available from: https://dx.plos.org/10.1371/journal.pone.0115571

28. Hong Z, Yang Y, Zhang C, Niu Y, Li K, Zhao X, et al. The retromer component SNX6 interacts with dynactin p150Glued and mediates endosome-to-TGN transport. Cell Res [Internet]. 2009 Dec [cited 2025 Mar 17];19(12):1334–49. Available from: https://www.nature.com/articles/cr2009130

29. Seaman MNJ. The retromer complex – endosomal protein recycling and beyond. Journal of Cell Science [Internet]. 2012 Jan 1 [cited 2025 Mar 17];jcs.103440. Available from: https://journals.biologists.com/jcs/article/doi/10.1242/jcs.103440/263210/The-retromer-complex-endosomal-protein-recycling

30. Liu J, Yin X, Liu B, Zheng H, Zhou G, Gong L, et al. HP1α mediates defective heterochromatin repair and accelerates senescence in Zmpste24-deficient cells. Cell Cycle. 2014;13(8):1237–47.

31. Gaillard H, García-Muse T, Aguilera A. Replication stress and cancer. Nat Rev Cancer [Internet]. 2015 May [cited 2023 Sep 17];15(5):276–89. Available from: https://www.nature.com/articles/nrc3916

32. Zeman MK, Cimprich KA. Causes and consequences of replication stress. Nat Cell Biol. 2014 Jan;16(1):2–9.

33. Nikolov I, Taddei A. Linking replication stress with heterochromatin formation. Chromosoma. 2016 Jun;125(3):523–33.

34. Gaggioli V, Lo CSY, Reverón-Gómez N, Jasencakova Z, Domenech H, Nguyen H, et al. Dynamic de novo heterochromatin assembly and disassembly at replication forks ensures fork stability. Nat Cell Biol [Internet]. 2023 Jul [cited 2023 Aug 19];25(7):1017–32. Available from: https://www.nature.com/articles/s41556-023-01167-z

35. Ayrapetov MK, Gursoy-Yuzugullu O, Xu C, Xu Y, Price BD. DNA double-strand breaks promote methylation of histone H3 on lysine 9 and transient formation of repressive chromatin. Proc Natl Acad Sci USA [Internet]. 2014 Jun 24 [cited 2023 Sep 13];111(25):9169–74. Available from: https://pnas.org/doi/full/10.1073/pnas.1403565111

36. Feng W, Jasin M. BRCA2 suppresses replication stress-induced mitotic and G1 abnormalities through homologous recombination. Nat Commun. 2017 Sep 13;8(1):525.

37. Hashimoto Y, Ray Chaudhuri A, Lopes M, Costanzo V. Rad51 protects nascent DNA from Mre11-dependent degradation and promotes continuous DNA synthesis. Nat Struct Mol Biol. 2010 Nov;17(11):1305–11.

38. Schlacher K, Christ N, Siaud N, Egashira A, Wu H, Jasin M. Double-strand break repair-independent role for BRCA2 in blocking stalled replication fork degradation by MRE11. Cell. 2011 May 13;145(4):529–42.

39. Lewis CM, Herbert BS, Bu D, Halloway S, Beck A, Shadeo A, et al. Telomerase immortalization of human mammary epithelial cells derived from a BRCA2 mutation carrier. Breast Cancer Res Treat. 2006 Sep;99(1):103–15.

40. Boutet-Robinet E, Trouche D, Canitrot Y. Neutral Comet Assay. BIO-PROTOCOL [Internet]. 2013 [cited 2024 Oct 25];3(18). Available from: https://bio-protocol.org/e915

41. Walsh KD, Kato TA. Alkaline Comet Assay to Detect DNA Damage. In: Gotoh E, editor. Chromosome Analysis [Internet]. New York, NY: Springer US; 2023 [cited 2024 Oct 25]. p. 65–72. (Methods in Molecular Biology; vol. 2519). Available from: https://link.springer.com/10.1007/978-1-0716-2433-3_7

42. Espinosa Jeffrey A, Blanchi B, Biancotti JC, Kumar S, Hirose M, Mandefro B, et al. Efficient Generation of Viral and Integration Free Human Induced Pluripotent Stem Cell Derived Oligodendrocytes. CP Stem Cell Biology [Internet]. 2016 Aug [cited 2024 Oct 25];38(1). Available from: https://currentprotocols.onlinelibrary.wiley.com/doi/10.1002/cpsc.11

43. Karaayvaz-Yildirim M, Silberman RE, Langenbucher A, Saladi SV, Ross KN, Zarcaro E, et al. Aneuploidy and a deregulated DNA damage response suggest haploinsufficiency in breast tissues of BRCA2 mutation carriers. Sci Adv. 2020 Jan;6(5):eaay2611.

44. Schlacher K, Wu H, Jasin M. A distinct replication fork protection pathway connects Fanconi anemia tumor suppressors to RAD51-BRCA1/2. Cancer Cell. 2012 Jul 10;22(1):106–16.

45. Kim JH, Na HK, Pak YK, Lee YS, Lee SJ, Moon A, et al. Roles of ERK and p38 mitogen-activated protein kinases in phorbol ester-induced NF-κB activation and COX-2 expression in human breast epithelial cells. Chemico-Biological Interactions [Internet]. 2008 Jan [cited 2025 Apr 16];171(2):133–41. Available from: https://linkinghub.elsevier.com/retrieve/pii/S0009279707002396

46. Canovas B, Nebreda AR. Diversity and versatility of p38 kinase signalling in health and disease. Nat Rev Mol Cell Biol [Internet]. 2021 May [cited 2022 Feb 2];22(5):346–66. Available from: http://www.nature.com/articles/s41580-020-00322-w

47. Löfmark S, de Klerk N, Aro H. Neisseria gonorrhoeae infection induces altered amphiregulin processing and release. PLoS One. 2011 Jan 27;6(1):e16369.

48. Piepkorn M, Underwood RA, Henneman C, Smith LT. Expression of amphiregulin is regulated in cultured human keratinocytes and in developing fetal skin. J Invest Dermatol. 1995 Dec;105(6):802–9.

49. Wang Y, Tzeng YDT, Chang G, Wang X, Chen S. Amphiregulin retains ERα expression in acquired aromatase inhibitor resistant breast cancer cells. Endocr Relat Cancer. 2020 Dec;27(12):671–83.

50. Panico L, D’Antonio A, Salvatore G, Mezza E, Tortora G, De Laurentiis M, et al. Differential immunohistochemical detection of transforming growth factor alpha, amphiregulin and CRIPTO in human normal and malignant breast tissues. Int J Cancer. 1996 Jan 3;65(1):51–6.

51. Yokoyama M, Ebert M, Funatomi H, Friess H, Buchler M, Johnson G, et al. AMPHIREGULIN IS A POTENT MITOGEN IN HUMAN PANCREATIC-CANCER CELLS – CORRELATION WITH PATIENT SURVIVAL. Int J Oncol [Internet]. 1995 Mar 1 [cited 2024 Oct 24]; Available from: http://www.spandidos-publications.com/10.3892/ijo.6.3.625

52. Hoogerbrugge N, Bult P, de Widt-Levert LM, Beex LV, Kiemeney LA, Ligtenberg MJL, et al. High prevalence of premalignant lesions in prophylactically removed breasts from women at hereditary risk for breast cancer. J Clin Oncol. 2003 Jan 1;21(1):41–5.

53. Kauff ND, Brogi E, Scheuer L, Pathak DR, Borgen PI, Hudis CA, et al. Epithelial lesions in prophylactic mastectomy specimens from women with BRCA mutations. Cancer. 2003 Apr 1;97(7):1601–8.

54. Cavalli V, Vilbois F, Corti M, Marcote MJ, Tamura K, Karin M, et al. The stress-induced MAP kinase p38 regulates endocytic trafficking via the GDI:Rab5 complex. Mol Cell. 2001 Feb;7(2):421–32.

55. Macé G, Miaczynska M, Zerial M, Nebreda AR. Phosphorylation of EEA1 by p38 MAP kinase regulates μ opioid receptor endocytosis. EMBO J [Internet]. 2005 Sep 21 [cited 2024 Oct 24];24(18):3235–46. Available from: http://emboj.embopress.org/cgi/doi/10.1038/sj.emboj.7600799

56. Sreekanth GP, Chuncharunee A, Sirimontaporn A, Panaampon J, Noisakran S, Yenchitsomanus P thai, et al. SB203580 Modulates p38 MAPK Signaling and Dengue Virus-Induced Liver Injury by Reducing MAPKAPK2, HSP27, and ATF2 Phosphorylation. Hsieh YH, editor. PLoS ONE [Internet]. 2016 Feb 22 [cited 2025 Mar 17];11(2):e0149486. Available from: https://dx.plos.org/10.1371/journal.pone.0149486

57. King D, Southgate HED, Roetschke S, Gravells P, Fields L, Watson JB, et al. Increased Replication Stress Determines ATR Inhibitor Sensitivity in Neuroblastoma Cells. Cancers (Basel). 2021 Dec 10;13(24):6215.

58. Moiseeva T, Hood B, Schamus S, O’Connor MJ, Conrads TP, Bakkenist CJ. ATR kinase inhibition induces unscheduled origin firing through a Cdc7-dependent association between GINS and And-1. Nat Commun [Internet]. 2017 Nov 9 [cited 2024 Oct 24];8(1):1392. Available from: https://www.nature.com/articles/s41467-017-01401-x

59. Raurell-Vila H, Bosch-Presegue L, Gonzalez J, Kane-Goldsmith N, Casal C, Brown JP, et al. An HP1 isoform-specific feedback mechanism regulates Suv39h1 activity under stress conditions. Epigenetics. 2017 Feb;12(2):166–75.

60. Maeda R, Tachibana M. HP1 maintains protein stability of H3K9 methyltransferases and demethylases. EMBO Rep. 2022 Apr 5;23(4):e53581.

61. Erdel F, Rademacher A, Vlijm R, Tünnermann J, Frank L, Weinmann R, et al. Mouse Heterochromatin Adopts Digital Compaction States without Showing Hallmarks of HP1-Driven Liquid-Liquid Phase Separation. Mol Cell. 2020 Apr 16;78(2):236–249.e7.

62. Goldman RD, Shumaker DK, Erdos MR, Eriksson M, Goldman AE, Gordon LB, et al. Accumulation of mutant lamin A causes progressive changes in nuclear architecture in Hutchinson–Gilford progeria syndrome. Proc Natl Acad Sci USA [Internet]. 2004 Jun 15 [cited 2024 Oct 24];101(24):8963–8. Available from: https://pnas.org/doi/full/10.1073/pnas.0402943101

63. De Sandre-Giovannoli A, Bernard R, Cau P, Navarro C, Amiel J, Boccaccio I, et al. Lamin A Truncation in Hutchinson-Gilford Progeria. Science [Internet]. 2003 Jun 27 [cited 2024 Oct 24];300(5628):2055–2055. Available from: https://www.science.org/doi/10.1126/science.1084125

64. Casasola A, Scalzo D, Nandakumar V, Halow J, Recillas-Targa F, Groudine M, et al. Prelamin A processing, accumulation and distribution in normal cells and laminopathy disorders. Nucleus. 2016;7(1):84–102.

65. Panzarino NJ, Krais JJ, Cong K, Peng M, Mosqueda M, Nayak SU, et al. Replication Gaps Underlie BRCA Deficiency and Therapy Response. Cancer Research [Internet]. 2021 Mar 1 [cited 2025 Apr 16];81(5):1388–97. Available from: https://aacrjournals.org/cancerres/article/81/5/1388/649177/Replication-Gaps-Underlie-BRCA-Deficiency-and

66. Tirman S, Quinet A, Wood M, Meroni A, Cybulla E, Jackson J, et al. Temporally distinct post-replicative repair mechanisms fill PRIMPOL-dependent ssDNA gaps in human cells. Molecular Cell [Internet]. 2021 Oct [cited 2025 Apr 16];81(19):4026–4040.e8. Available from: https://linkinghub.elsevier.com/retrieve/pii/S1097276521007474

67. De Feraudy S, Revet I, Bezrookove V, Feeney L, Cleaver JE. A minority of foci or pan-nuclear apoptotic staining of γH2AX in the S phase after UV damage contain DNA double-strand breaks. Proc Natl Acad Sci USA [Internet]. 2010 Apr 13 [cited 2025 May 12];107(15):6870–5. Available from: https://pnas.org/doi/full/10.1073/pnas.1002175107

68. Sluss HK, Davis RJ. H2AX Is a Target of the JNK Signaling Pathway that Is Required For Apoptotic DNA Fragmentation. Molecular Cell [Internet]. 2006 Jul [cited 2025 May 12];23(2):152–3. Available from: https://linkinghub.elsevier.com/retrieve/pii/S1097276506004576

69. Baure J, Izadi A, Suarez V, Giedzinski E, Cleaver JE, Fike JR, et al. Histone H2AX phosphorylation in response to changes in chromatin structure induced by altered osmolarity. Mutagenesis. 2009 Mar;24(2):161–7.

70. Liu W. Single Molecular Resolution to Monitor DNA Replication Fork Dynamics upon Stress by DNA Fiber Assay. Bio Protoc. 2021 Dec 20;11(24):e4269.

71. Blow JJ, Ge X. Replication forks, chromatin loops and dormant replication origins. Genome Biol [Internet]. 2008 [cited 2024 Oct 24];9(12):244. Available from: http://genomebiology.biomedcentral.com/articles/10.1186/gb-2008-9-12-244

72. Fu H, Baris A, Aladjem MI. Replication timing and nuclear structure. Curr Opin Cell Biol. 2018 Jun;52:43–50.

73. Kotsantis P, Silva LM, Irmscher S, Jones RM, Folkes L, Gromak N, et al. Increased global transcription activity as a mechanism of replication stress in cancer. Nat Commun. 2016 Oct 11;7:13087.

74. Dworak N, Makosa D, Chatterjee M, Jividen K, Yang CS, Snow C, et al. A nuclear lamina-chromatin-Ran GTPase axis modulates nuclear import and DNA damage signaling. Aging Cell. 2019 Feb;18(1):e12851.

75. Her J, Ray C, Altshuler J, Zheng H, Bunting SF. 53BP1 Mediates ATR-Chk1 Signaling and Protects Replication Forks under Conditions of Replication Stress. Mol Cell Biol. 2018 Apr 15;38(8):e00472–17.

76. Moudry P, Lukas C, Macurek L, Neumann B, Heriche JK, Pepperkok R, et al. Nucleoporin NUP153 guards genome integrity by promoting nuclear import of 53BP1. Cell Death Differ [Internet]. 2012 May [cited 2024 Oct 24];19(5):798–807. Available from: https://www.nature.com/articles/cdd2011150

77. Galanty Y, Belotserkovskaya R, Coates J, Polo S, Miller KM, Jackson SP. Mammalian SUMO E3-ligases PIAS1 and PIAS4 promote responses to DNA double-strand breaks. Nature. 2009 Dec 17;462(7275):935–9.

78. Shevelyov YY. Interactions of Chromatin with the Nuclear Lamina and Nuclear Pore Complexes. Int J Mol Sci. 2023 Oct 30;24(21):15771.

79. Bozler J, Nguyen HQ, Rogers GC, Bosco G. Condensins Exert Force on Chromatin-Nuclear Envelope Tethers to Mediate Nucleoplasmic Reticulum Formation in *Drosophila melanogaster*. G3 Genes|Genomes|Genetics [Internet]. 2015 Mar 1 [cited 2024 Oct 25];5(3):341–52. Available from: https://academic.oup.com/g3journal/article/5/3/341/6058727

80. Stephens AD, Liu PZ, Banigan EJ, Almassalha LM, Backman V, Adam SA, et al. Chromatin histone modifications and rigidity affect nuclear morphology independent of lamins. Mol Biol Cell. 2018 Jan 15;29(2):220–33.

81. Stiekema M, Houben F, Verheyen F, Borgers M, Menzel J, Meschkat M, et al. The Role of Lamins in the Nucleoplasmic Reticulum, a Pleiomorphic Organelle That Enhances Nucleo-Cytoplasmic Interplay. Front Cell Dev Biol [Internet]. 2022 Jun 16 [cited 2024 Oct 24];10:914286. Available from: https://www.frontiersin.org/articles/10.3389/fcell.2022.914286/full

82. Rusiñol AE, Sinensky MS. Farnesylated lamins, progeroid syndromes and farnesyl transferase inhibitors. Journal of Cell Science [Internet]. 2006 Aug 15 [cited 2025 Jan 2];119(16):3265–72. Available from: https://journals.biologists.com/jcs/article/119/16/3265/29042/Farnesylated-lamins-progeroid-syndromes-and

83. Goulbourne CN, Malhas AN, Vaux DJ. The induction of a nucleoplasmic reticulum by prelamin A accumulation requires CTP:phosphocholine cytidylyltransferase-α. J Cell Sci. 2011 Dec 15;124(Pt 24):4253–66.

84. Chan KL, Palmai-Pallag T, Ying S, Hickson ID. Replication stress induces sister-chromatid bridging at fragile site loci in mitosis. Nat Cell Biol. 2009 Jun;11(6):753–60.

85. Vincent J, Adura C, Gao P, Luz A, Lama L, Asano Y, et al. Small molecule inhibition of cGAS reduces interferon expression in primary macrophages from autoimmune mice. Nat Commun. 2017 Sep 29;8(1):750.

86. Mackenzie KJ, Carroll P, Martin CA, Murina O, Fluteau A, Simpson DJ, et al. cGAS surveillance of micronuclei links genome instability to innate immunity. Nature. 2017 Aug 24;548(7668):461–5.

87. Yang H, Wang H, Ren J, Chen Q, Chen ZJ. cGAS is essential for cellular senescence. Proc Natl Acad Sci U S A. 2017 Jun 6;114(23):E4612–20.

88. Crossley MP, Song C, Bocek MJ, Choi JH, Kousorous J, Sathirachinda A, et al. R-loop-derived cytoplasmic RNA–DNA hybrids activate an immune response. Nature [Internet]. 2023 Jan 5 [cited 2023 Mar 4];613(7942):187–94. Available from: https://www.nature.com/articles/s41586-022-05545-9

89. Hansen AM, Ge Y, Schuster MB, Pundhir S, Jakobsen JS, Kalvisa A, et al. H3K9 dimethylation safeguards cancer cells against activation of the interferon pathway. Sci Adv. 2022 Mar 18;8(11):eabf8627.

90. Allshire RC, Madhani HD. Ten principles of heterochromatin formation and function. Nat Rev Mol Cell Biol [Internet]. 2018 Apr [cited 2024 Oct 24];19(4):229–44. Available from: https://www.nature.com/articles/nrm.2017.119

91. Barrowman J, Hamblet C, George CM, Michaelis S. Analysis of prelamin A biogenesis reveals the nucleus to be a CaaX processing compartment. Mol Biol Cell. 2008 Dec;19(12):5398–408.

92. Lattanzi G, Columbaro M, Mattioli E, Cenni V, Camozzi D, Wehnert M, et al. Pre-Lamin A processing is linked to heterochromatin organization. J Cell Biochem. 2007 Dec 1;102(5):1149–59.

93. Chaturvedi P, Parnaik VK. Lamin A rod domain mutants target heterochromatin protein 1alpha and beta for proteasomal degradation by activation of F-box protein, FBXW10. PLoS One. 2010 May 13;5(5):e10620.

94. Baldeyron C, Soria G, Roche D, Cook AJL, Almouzni G. HP1alpha recruitment to DNA damage by p150CAF-1 promotes homologous recombination repair. J Cell Biol. 2011 Apr 4;193(1):81–95.

95. Ogawa K, Utsunomiya T, Mimori K, Tanaka F, Haraguchi N, Inoue H, et al. Differential gene expression profiles of radioresistant pancreatic cancer cell lines established by fractionated irradiation. Int J Oncol [Internet]. 2006 Mar 1 [cited 2025 Apr 16]; Available from: http://www.spandidos-publications.com/10.3892/ijo.28.3.705

96. Eckstein N, Servan K, Girard L, Cai D, von Jonquieres G, Jaehde U, et al. Epidermal growth factor receptor pathway analysis identifies amphiregulin as a key factor for cisplatin resistance of human breast cancer cells. J Biol Chem. 2008 Jan 11;283(2):739–50.

97. Yoshida M, Shimura T, Fukuda S, Mizoshita T, Tanida S, Kataoka H, et al. Nuclear translocation of pro-amphiregulin induces chemoresistance in gastric cancer. Cancer Sci. 2012 Apr;103(4):708–15.

98. Soufi A, Donahue G, Zaret KS. Facilitators and impediments of the pluripotency reprogramming factors’ initial engagement with the genome. Cell. 2012 Nov 21;151(5):994–1004.

99. Lim E, Wu D, Pal B, Bouras T, Asselin-Labat ML, Vaillant F, et al. Transcriptome analyses of mouse and human mammary cell subpopulations reveal multiple conserved genes and pathways. Breast Cancer Res. 2010;12(2):R21.

100. Tollot-Wegner M, Jessen M, Kim K, Sanz-Moreno A, Spielmann N, Gailus-Durner V, et al. TRPS1 maintains luminal progenitors in the mammary gland by repressing SRF/MRTF activity. Breast Cancer Res [Internet]. 2024 May 3 [cited 2025 Feb 7];26(1):74. Available from: https://breast-cancer-research.biomedcentral.com/articles/10.1186/s13058-024-01824-7

101. Macheret M, Halazonetis TD. DNA replication stress as a hallmark of cancer. Annu Rev Pathol. 2015;10:425–48.

