## Supplementary Methods for "The EGFR ligand amphiregulin regulates genomic integrity by facilitating heterochromatin formation in response to replication stress"

**Jiang et al. Supplementary Methods:**

Antibody list

qRT-PCR

Immunofluorescence/Immunohistochemistry

Senescence associated beta-galactosidase detection

Mice

Patient samples

Microscopy

**Antibodies**

| **Primary antibodies** | **Source** | **Catalogue #** |
|---|---|---|
| 53BP1 | Novus Biologicals | NB100304 |
| Actin | Sigma | A-2066 |
| ATM | Calbiochem | PC116 |
| Amphiregulin | Proteintech | AF262 |
| Amphiregulin | Proteintech | 16036-1-AP |
| Amphiregulin | Santa Cruz Biotech | H-155: sc-25436 |
| BRCA2 | Abcam | ab27976 |
| BrdU [MoBu-1] | Abcam | ab8039 |
| BU1/75 [ICR1] | Abcam | ab6326 |
| BrdU [B44] | Becton Dickson | 347580 (7580) |
| cGAS | Cell Signaling | 79978 |
| CHK1 | Cell Signaling | 2360 |
| CHK2 (1C12) | Cell Signaling | 3440 |
| DDX52 | Bethyl Laboratories | A303-053A |
| EEA1 | Cell Signaling | 3288 |
| GAPDH | Cell Signaling | 97166S |
| gH2AX | Cell Signaling | 2577 |
| H3K9me3 | Thermo | PA5-31910 |
| HP1α | Sigma | 05-689 |
| Lamin A/C | SantaCruz | sc-7292 |
| Lamin A/C | Cell Signaling | 2032 |
| Mab 414 | Biolegend | 902901 |
| P-ser1981 ATM | Calbiochem | DR1002 |
| p38 MAPK | Cell Signaling | 9212 |
| phospho-p38 | Cell Signaling | D3F9 |
| P-thr68 CHK2 | Thermo | PA5-17818 |
| P-ser345 CHK1 | Cell Signaling | 2348 |
| Prelamin A (PL-1C7) | Millipore Sigma | MABT858 |
| RAN | Millipore | 07-517 |
| SUV39h1 | ProteinTech | 10574-1-AP |
| Vinculin | Abcam | ab129002 |
| V5-Tag | Abcam | AB27671 |
| **Secondary antibodies** | **Source** | **Catalogue#** |
| Rabbit IgG (HRP-linked) | Cell Signaling | 7074S |
| Mouse IgG (HRP-linked) | Cell Signaling | 7076 |
| Goat IgG (HRP-linked) | Invitrogen | A15999 |

**qRT-PCR:** Cells were lysed after culture and the RNA is extracted using the Aurum™ Total RNA mini kit (BIO-RAD, 732-6820) following the manufacturer's protocol and was resuspended in 40ul of nuclease-free water and stored at -80°C. cDNA was synthesized by reverse transcribing 1ug of RNA with reverse transcriptase (BIO-RAD, 1708891). The resulting cDNA was diluted to 20ng/ul and mixed at 1:10 ratio with the SsoAdvanced Universal SYBR Green Supermix (BIO-RAD, 1725271). The qRT-PCR reactions were performed using the CFX96 Real-Time PCR System [C-1000 Touch Thermal Cycler] (BIO-RAD) and relative gene expressions were calculated using the $\Delta\Delta C_T$ method normalized to S18 housekeeping gene. Primers were designed online using the National Center for Biotechnology Information (https://www.ncbi.nlm.nih.gov/tools/primer-blast/) and manufactured by Invitrogen/ Thermo Fisher. Primer sequences can be found in Supplementary Table 1.

**Lentivirus production:** HEK293T cells were seeded into 10 cm plates and 3 days later the plasmid transfection mix was prepared by combining 240 µL Optimem, 8 µg transfer plasmids, 4 µg PCMV (envelope plasmid), 6 µg pMD2.G (packaging plasmid), and 72 µL PEI (4 µL of PEI/1 µg DNA). The DNA:PEI mixture was gently mixed for 20 minutes at room temperature, then diluted to a total volume of 3 mL with Optimem media. The mixture was then added onto cells. The following day (D4), Optimem was removed and replaced with 10 mL of fresh media (DMEM +10%FBS). At 48h and 72h post infection, media containing virus were obtained and stored at 4°C. A virus precipitation kit (Benchmark bioscience) was used to concentrate and store the lentivirus.

**Immunofluorescence/Immunohistochemistry:** Immunofluorescence was performed with paraffin sections and fixed hTERT-MECs. Cells were fixed with 4% PFA for 15mins at room temperature whereas paraffin sections were pre-fixed and mounted onto microscope slides. Both type of samples were incubated with a blocking solution (5% BSA, 0.5% Triton-X in 1X PBS) at room temperature for 1 hour. Antibodies were diluted using dilution buffer (1% BSA. 0.1% Triton-X in 1X PBS). Samples were incubated with primary antibodies at 4°C overnight followed by 3 × 5 min PBS wash. Sections were then incubated with secondary antibodies at RT for 1h. Subsequently, the samples were washed again with 1X PBS and counterstained with DAPI (Thermo Fisher Scientific, R37606) for 10 mins before mounting with antifade mounting media (Vector laboratory, Vectashield H1900).

Immunohistochemistry was performed using paraffin sections. After the slides were deparaffinized, they were incubated with 3% H2O2 at room temperature for 20 mins. After 3 x 5 mins PBS wash, the slides were incubated with a blocking solution (DAKO, X0909). Antibodies were diluted using dilution buffer (DAKO, S0809) to working concentrations and applied for overnight incubation at 4°C. The slides were washed 3 × 5 min PBS the following day and HRP-anti-rabbit was applied onto the samples and incubated at room temperature for 1h. After Incubation, DAB working solution was prepared (DAKO, K346711). Application of DAB was time sensitive and a close monitoring under the microscope was needed for the development of a brown colour. Subsequently, the slides were washed and counterstained with hematoxylin (Abcam, ab245880). Dehydration was performed in 70%, 95% and 100% ethanol followed by toluene. Finally, the slides were mounted with coverslips using permanent mounting media (Permount, ThermoFisher).

**Senescence associated β-galactosidase detection:** Detection of SAβ-gal was performed as described by Itahana et al (1) for cultured cells.

**Western blot:** Cells were lysed using RIPA buffer (50 mM Tris-HCl, pH 7.4, 0.1% sodium deoxycholate, 1 mM EDTA, 140 mM NaCl, 1% Triton-X, 0.1% SDS), and total cell lysates were collected prior to the addition of Laemmli buffer (Bio-Rad, #1610747). Protein samples were denatured at 95°C for 5 minutes before running on an 8–12% acrylamide gel. Proteins were transferred to a PVDF membrane for subsequent probing. Specific proteins were detected using the appropriate primary antibodies and horseradish peroxidase-conjugated secondary antibodies (see antibody list). Protein bands were visualized using the Chemiluminescent Substrate ECL (Thermo Fisher Scientific) and imaged on a ChemiDoc MP Imaging System (Bio-Rad).

**Mice:** FVB/n *Brca2*$^{flox}$ mice (Jackson Labs JAX stock #032752) were bred with FVB/n MMTV-cre mice (the gift of Dr. William Muller) to produce *Brca2*+/+, +/-, -/- genotypes. Euthansia is performed subsequently to describe treatment and mammary glands were harvested for paraffin embedding and RNA extraction. Genotyping of FVB/N MMTV-cre; *Brca2* floxed mice ear punch DNA was performed using forward and reverse primers: GGCTGTCTTAGAACTTAGGCTG and CTCCACACATACATCATGTGTC respectively for *Brca2*, GGGATTGCTTATAACACCCTGTTACG and TATTCGGATCATCAGCTACACCAGAG respectively for Cre. Animals were housed on a protocol approved by the University of Ottawa

Animal Care and Veterinary Services Committee and in accordance with Canadian Council on Animal Care guidelines. Mammary glands were isolated immediately after euthanasia. Samples were fixed in 10% formalin at 4°C for 24h. Fixed samples were transferred in 70% EtOH to University of Ottawa Department of Pathology and Laboratory Medicine for paraffin embedding and sectioning.

**Patient samples:** Human breast tissue samples were harvested from prophylactic mastectomy tissue from patients harbouring a germline BRCA2 mutation confirmed by the Children's Hospital of Eastern Ontario Genetics Diagnostics Lab. Non-carrier tissue was obtained from reduction mammoplasty surgery. Tissue collection was approved by REB with the Ottawa Hospital Ethics Board. Tissues were collected at surgery in collection media (DMEM/F12) and fixed in 10% formalin at 4°C for 72h then transferred in 70% EtOH and delivered to University of Ottawa Department of Pathology and Laboratory Medicine for paraffin embedding and sectioning.

**Microscopy:** Fluorescence imaging was performed with a Zeiss AxioObserver 7-APOllo inverted epifluorescence microscope using a Plan-Apochromat ×63/1.4 oil objective. Images were captured using a Hamamatsu ORCA-Flash LT sCMOS (monochrome) camera and ZEN 3.1 pro software or using a Zeiss Axiocam 705 Colour camera. Imaging of IHC was performed with a Thermo Fisher FL Auto 2 (inverted) microscope using 20X, 0.4 NA air objective for general sample visualization. Images were captured using a colour camera and Auto2 software. Confocal images were acquired using a Zeiss AxioObserverZ1 mot inverted confocal microscope using a 63x, 1.4 NA, oil, Plan-Apo objective.

1. Itahana K, Campisi J, Dimri GP. Methods to detect biomarkers of cellular senescence: the senescence-associated beta-galactosidase assay. Methods Mol Biol. 2007;371:21–31.
